# A kinetic model of the central carbon metabolism for acrylic acid production in *Escherichia coli*

**DOI:** 10.1101/2020.05.13.093294

**Authors:** Alexandre Oliveira, Joana Rodrigues, Eugénio Ferreira, Lígia Rodrigues, Oscar Dias

**Affiliations:** Centre of Biological Engineering, University of Minho, Braga, Portugal

## Abstract

Acrylic acid is a value-added chemical used in industry to produce diapers, coatings, paints, and adhesives, among many others. Due to its economic importance, there is currently a need for new and sustainable ways to synthesise it. Recently, the focus has been laid in the use of *Escherichia coli* to express the full bio-based pathway using 3-hydroxypropionate as an intermediary through three distinct pathways (glycerol, malonyl-CoA, and *β*-alanine). Hence, the goals of this work were to use COPASI software to assess which of the three pathways has a higher potential for industrial-scale production, from either glucose or glycerol, and identify potential targets to improve the biosynthetic pathways yields.

When compared to the available literature, the models developed during this work successfully predict the production of 3-hydroxypropionate, using glycerol as carbon source in the glycerol pathway, and using glucose as a carbon source in the malonyl-CoA and *β*-alanine pathways. Finally, this work allowed to identify four potential over-expression targets (glycerol-3-phosphate dehydrogenase (G3pD), acetyl-CoA carboxylase (AccC), aspartate aminotransferase (AspAT), and aspartate carboxylase (AspC)) that should, theoretically, result in higher AA yields.

**Author summary:** Acrylic acid is an economically important chemical compound due to its high market value. Nevertheless, the majority of acrylic acid consumed worldwide its produced from petroleum derivatives by a purely chemical process, which is not only expensive, but it also contributes towards environment deterioration. Hence, justifying the current need for sustainable novel production methods that allow higher profit margins. Ideally, to minimise production cust, the pathway should consist in the direct bio-based production from microbial feedstocks, such as Escherichia coli, but the current yields achieved are still to low to compete with conventional method. In this work, even though the glycerol pathway presented higher yields, we identified the malonyl-CoA route, when using glucose as carbon source, as having the most potential for industrial-scale production, since it is cheaper to implement. Furthermore, we also identified potential optimisation targets for all the tested pathways, that can help the bio-based method to compete with the conventional process.

## Introduction

Acrylic acid (AA) (C_3_H_4_O_2_) is an important chemical compound that is one of the key components of superabsorbent polymers [1–3]. According to the Allied Market Research, in 2015, the global market for AA was valued at 12,500 million US dollars, and is expected to reach 19,500 million US dollars until 2022 [4]. Despite its economic importance, the vast majority of AA is still produced by the oxidation of propylene or propane in a purely chemical process [1,5,6]. Ergo, the principal method for AA production was found to be expensive, with a high energy demand, thus contributing to the planet’s environment decay. Hence, the development of an innovative and sustainable biological production method has been attracting the attention of the scientific community [1,2,7]. In the last decade, several semi-biological methods have emerged and were optimised. These methods consist of the bio-based production of 3-hydroxypropionate (3-HP) and its subsequent chemical conversion to AA. Despite the substantial improvements obtained with these methods, this process involves a catalytic step that increases the production costs and environmental impact due to high energy demands [1–3,5]. Hence, the AA’s production method should, ideally, be a bio-based direct route as, in theory, microbial feedstocks are less expensive, allowing a higher profit margin [1]. Moreover, a more sustainable bioprocess allows to decrease non-renewable resources dependence and CO_2_ emissions.

Fortunately, in recent years, it has been proven that it is possible to use engineered *Escherichia coli* to convert glucose or glycerol into AA. Like in the semi-biological methods, the bioprocess is also divided into two main parts, the production of 3-HP and its subsequent conversion to AA. This part of the pathway, from 3-HP to AA, has not been extensively studied. So far, there are only three studies that successfully converted glucose or glycerol to AA in *E. coli* [1,2,7]. However, the synthesis of 3-HP is well reported, and three distinct pathways have been identified, namely the glycerol route, the malonyl-CoA route, and the *β*-alanine route. From these pathways, it is well established that the glycerol pathway is associated with the highest yields. However, one of the reactions of this route requires the supplementation of vitamin B_12_ (Fig 1), which is an expensive practice at an industrial-scale production, hence a significant disadvantage of this route [8,9].

**Fig 1.**
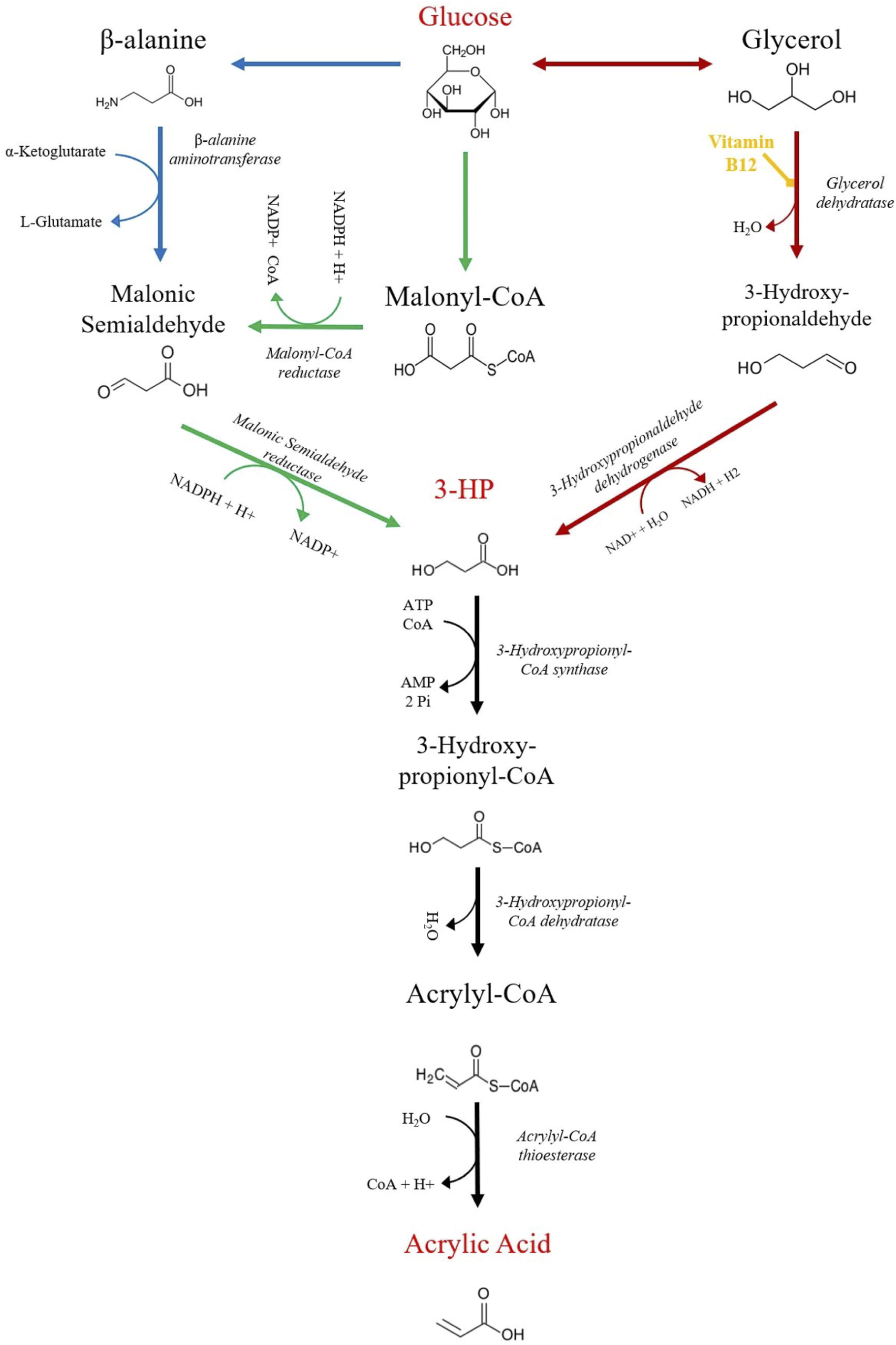
Biosynthetic pathways for acrylic acid (AA) production from Glucose using 3-hydroxypropionate (3-HP) as an intermediary. 3-HP can be produced from Glucose through three distinct pathways: glycerol (red arrows), malonyl-CoA (green arrows), and *β*-alanine (blue arrows). Furthermore, *E. coli* can also direct Glycerol towards the central carbon metabolism, allowing it to be used as a carbon source.

The bio-based method is currently considered a promising alternative to the conventional process as the production of 3-HP increased considerably in the last few years. Recently, studies reported productions of up to 8.10 g/L with the glycerol pathway [1], 3.60 g/L with the malonyl-CoA pathway [10], and 0.09 g/L with the *β*-alanine pathway [11]. However, the AA yields obtained by Tong et al. (2016) [2] (0.0377 g/L) and Chu et al. (2015) [1] (0.12 g/L) for the glycerol pathway, and Liu and Liu (2016) [7] (0.013 g/L) for the malonyl-CoA pathway, established that this process still needs to be optimised to compete with the currently used methods.

Taking these considerations into account, the main goals of this work are to identify the reactions of the known routes for AA production (glycerol, malonyl-CoA, and β-alanine pathways) and to determine which pathway have a higher potential for industrial-scale production. *E. coli*’s central carbon metabolism (CCM) kinetic models will be used to analyse the three pathways using either glucose or glycerol as carbon source. Finally, novel optimisation strategies to improve the AA yields of the three biosynthetic pathways will also be sought.

## Results and Discussion

### Time Course Simulations

During the simulations, several issues arose, leading to changes in parameters before the analysis of the 3-HP and AA production. These changes are explained in detail in the S1 Appendix, section 1.1.

Regarding the production of 3-HP in the glycerol pathway, simulations with models set to use either glucose (Glu-Gly) or glycerol as carbon source (Gly-Gly), predicted, the production of 0.19 g/L (after three hours), and 8.30 g/L (after six-hours), respectively (Fig 3). Whereas, concerning the production of AA, the Glu-Gly and Gly-Gly models predicted 0.16 g/L and 6.71 g/L, respectively (Fig 4). From these results, glycerol seems to be associated with higher yields, which is in good agreement with the available literature [1]. Moreover, regarding the production of AA, the intracellular concentration of 3-HP showed that there is no accumulation (Fig 4), meaning that most 3-HP is converted into AA. These results are most likely associated with the use of excessive enzyme concentration to calculate the V_max_ for the heterologous pathway, which led to a state in which the main limiting factor in the synthesis of AA was the CCM’s flux distribution. However, this is not the case *in vivo*, as the studies that tested the full bio-based pathway show that 3-HP and other intermediates indeed accumulate during this process [1,2].

**Fig 2.**
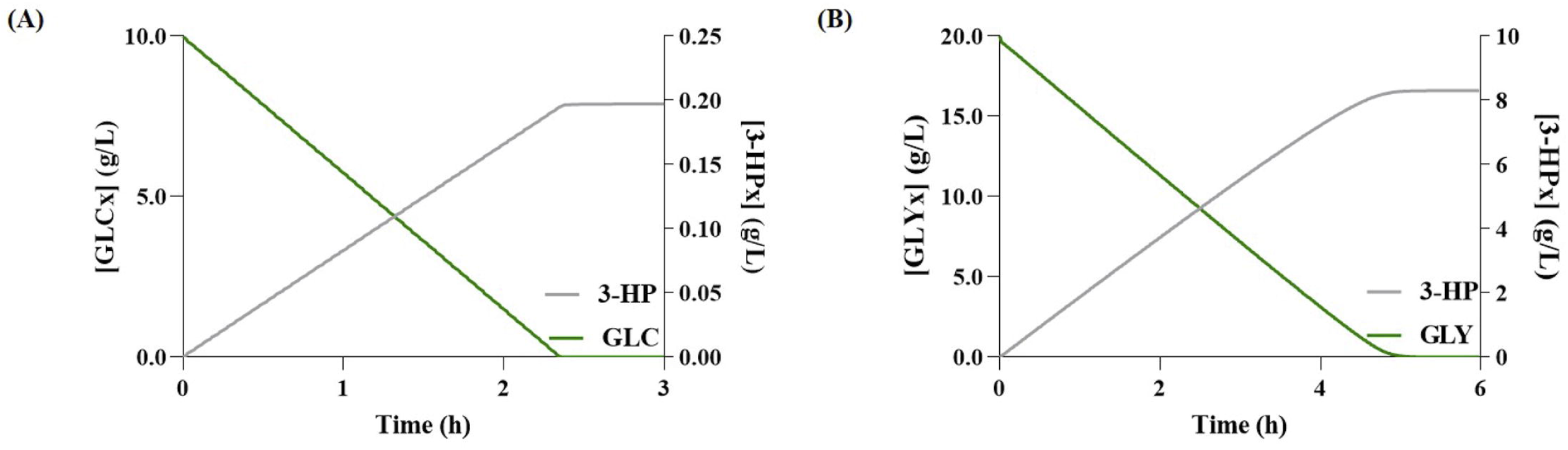
Simulation results for 3-hydroxypropionate (3-HP) production via the glycerol pathway. **(A)** Glucose (GLCx) consumption and variation of extracellular 3-HP (3-HPx) over time; **(B)** Glycerol (GLYx) consumption and variation of 3-HPx over time.

**Fig 3.**
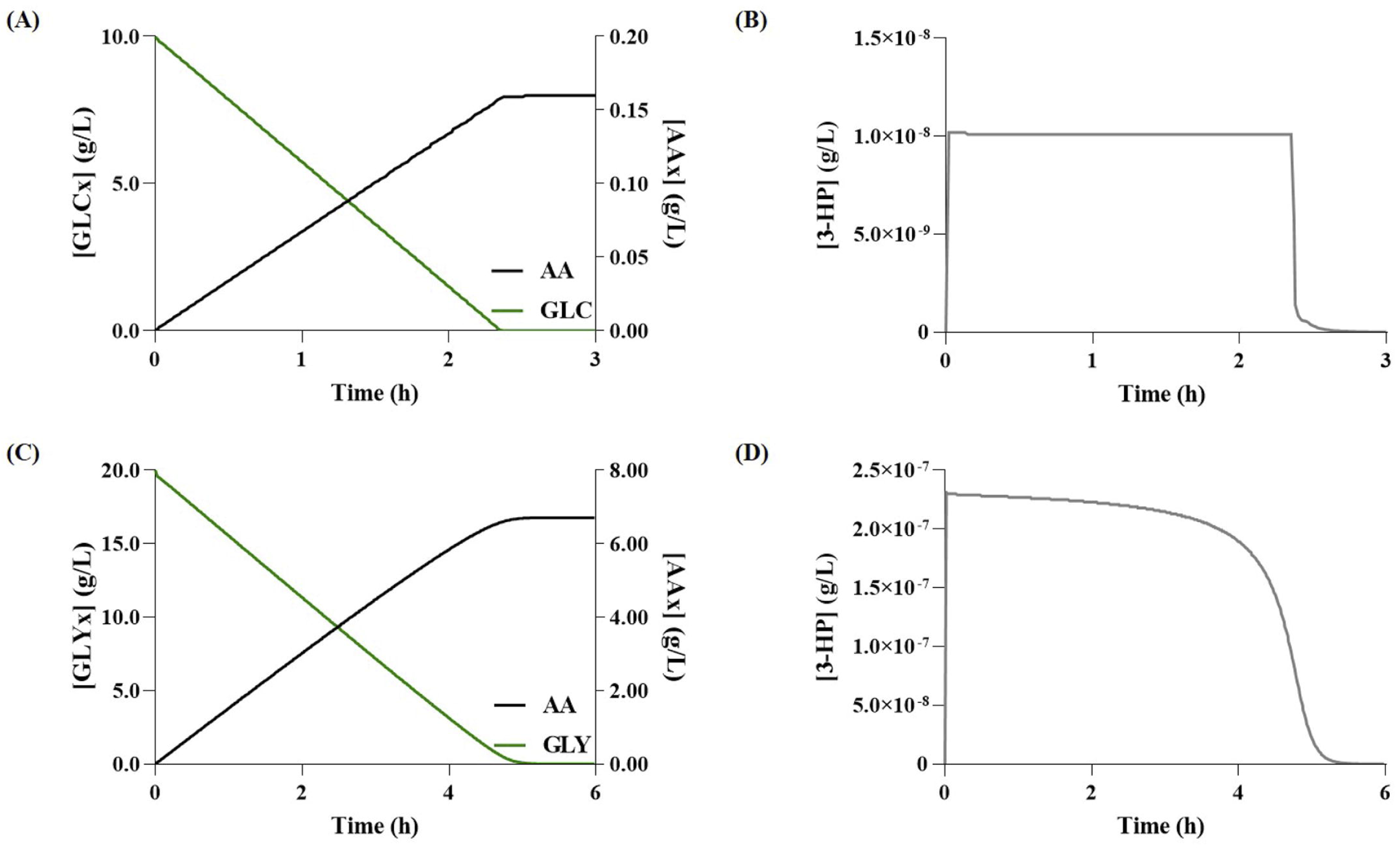
Simulation results for acrylic acid (AA) production via the glycerol pathway. **(A)** Glucose (GLC) consumption and variation of extracellular AA (AAx) over time; **(B)** Variation of 3-hydroxypropionate (3-HP) concentration over time when using Glucose as carbon source; **(C)** Glycerol (GLYx) consumption and variation of extracellular AAx over time; **(D)** Variation of 3-HP concentration over time when using Glycerol as carbon source.

**Fig 4.**
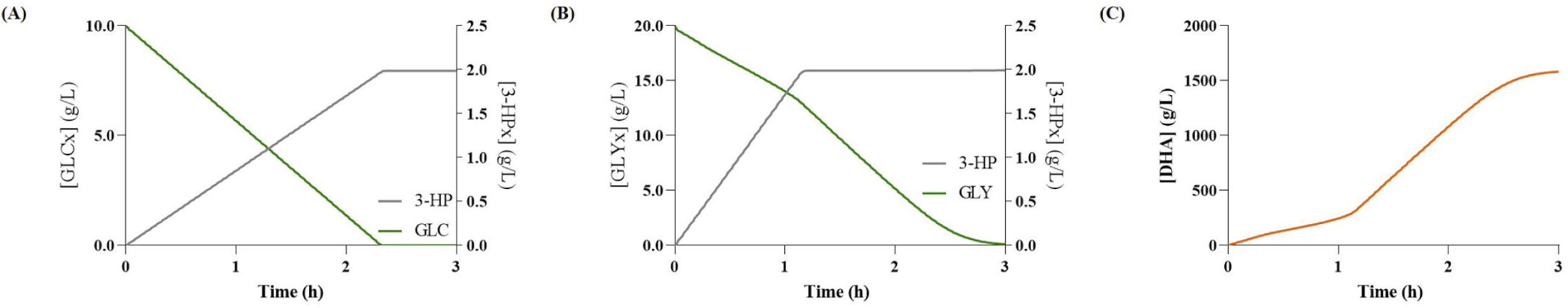
Simulation results for 3-hydroxypropionate (3-HP) production via the malonyl-CoA pathway. **(A)** Glucose (GLCx) consumption and variation of extracellular 3-HP (3-HPx) over time; Glycerol (GLYx) consumption and variation of 3-HPx over time; **(C)** Variation of intracellular dihydroxyacetone (DHA) concentration over time when using Glycerol.

When comparing the predictions of the glycerol pathway models (Table 3), it is possible to observe that the predicted 3-HP concentration, with the Gly-Gly model, is slightly different from literature reports. This difference slightly increases when increasing the initial concentration of carbon. For instance, for 40 g/L of glycerol, the predicted 3-HP production is about two times higher. Nevertheless, the model representing the 3-HP production from glycerol exhibits promising results.

**Table 1.** Literature review on 3-hydroxypropionate (3-HP) and acrylic acid (AA) production yields by the glycerol pathway in metabolically engineered *Escherichia coli*, and comparison with the yields predicted by the dynamic models using the same initial carbon concentration.

| Reference | End Product | Carbon Source | Initial Carbon Conc. (g/L) | Titer (g/L) | Predicted Titer (g/L) |
| --- | --- | --- | --- | --- | --- |
| Raj et al. (2009)[12] | 3-HP | Glycerol | 9.20 | 2.80 | 3.59 |
| Rathnasingh et al. (2009) [13] | 3-HP | Glycerol | 18.40 | 4.40 | 7.59 |
| Chu et al. (2015) [14] | 3-HP | Glycerol | 40.00 | 7.40 | 17.19 |
| Chu et al. (2015) [1] | 3-HP | Glycerol | 40.00 | 8.10 | 17.19 |
|  |  | Glucose | 21.50 | 3.90 | 0.42 |
| Tong et al. (2016) [2] | AA | Glycerol | 20.00 | 0.037 | 8.30 |

**Table 2.** Literature review on 3-hydroxypropionate (3-HP) and acrylic acid (AA) production yields by the malonyl-CoA pathway in metabolically engineered *Escherichia coli*, and comparison with the yields predicted by the dynamic models using the same initial carbon concentration.

| Reference | End Product | Carbon Source | Initial Carbon Conc. (g/L) | Titer (g/L) | Predicted Titer (g/L) |
| --- | --- | --- | --- | --- | --- |
| Cheng et al. (2016) [16] | 3-HP | Glucose | 10.00 | 1.80 | 1.99 |
| Liu et al. (2016) [10] | 3-HP | Glucose | 20.00 | 3.60 | 3.96 |
| Liu and Liu (2016) [7] | AA | Glucose | 20.00 | 0.013 | 3.22 |

**Table 3.** Literature review on 3-hydroxypropionate (3-HP) production yields by the *β*-alanine pathway in metabolically engineered *Escherichia coli*, and comparison with the yields predicted by the dynamic models using the same initial carbon concentration.

| Reference | End Product | Carbon Source | Initial Carbon Conc. (g/L) | Titer (g/L) | Predicted Titer (g/L) |
| --- | --- | --- | --- | --- | --- |
| Song et al. (2016) [11] | 3-HP | Glucose | 15.00 | 0.09 | 0.049 |

The scenario with the Glu-Gly model is considerably distinct, as a ten-fold lower concentration of 3-HP was predicted. This difference might be associated with the production of glycerol, more specifically, in the flux through G3pD and G3pP, as according to Chu et al. (2015) [1] their strain is able to accumulate more glycerol (2.5 g/L) than this model is able to produce (0.34 g/L), for the same amount of glucose. Unfortunately, such study, which presented the highest AA concentration (0.12 g/L) reported thus far, did not disclose the amount of glucose used to obtain such production. Hence, it was not possible to directly compare the predicted titer from the Glu-Gly model.

When performing simulation using glucose concentrations of 10 g/L and 20 g/L, the model predicts the production of 0.16 g/L and 0.32 g/L of AA, respectively. These results show that the AA production predicted by our model would be in good agreement with the results reported by Chu et al. (2015) [1], if 10 g/L of glucose had been used for the carbon source. However, the model does not accumulate any intermediary compounds of the heterologous pathway; thus, most 3-HP is converted into AA, which does not correctly represent the *in vivo* results. Consequently, when assessing the results to the work of Tong et al. (2016) [2] that tested the production of AA in *E. coli*, the model fails to predict the AA yield accurately, as expected.

Regarding the malonyl-CoA pathway, the model set to use glucose as carbon source (Glu-Mcoa) predicted a titer of 1.99 g/L of 3-HP, while the malonyl-CoA model set to use glycerol as carbon source (Gly-Mcoa) predicted 1.99 g/L of 3-HP (Fig 5). Moreover, the Glu-Mcoa and Gly-Mcoa models predicted the production of 1.62 g/L and 0.17 g/L of AA, respectively (Fig 6). The behaviour analysis of the model showed a considerable intracellular accumulation of dihydroxyacetone in the Gly-Mcoa model (Fig 5C). Hence, most carbon does not reach the CCM, and therefore these results should not be considered as it is not possible to determine the best carbon source to produce AA. Although literature reports suggest a consensus towards the use of glucose as a carbon source, it should be noted that no work using glycerol was found. Thus, glycerol should not be excluded as a promising alternative carbon source.

**Fig 5.**
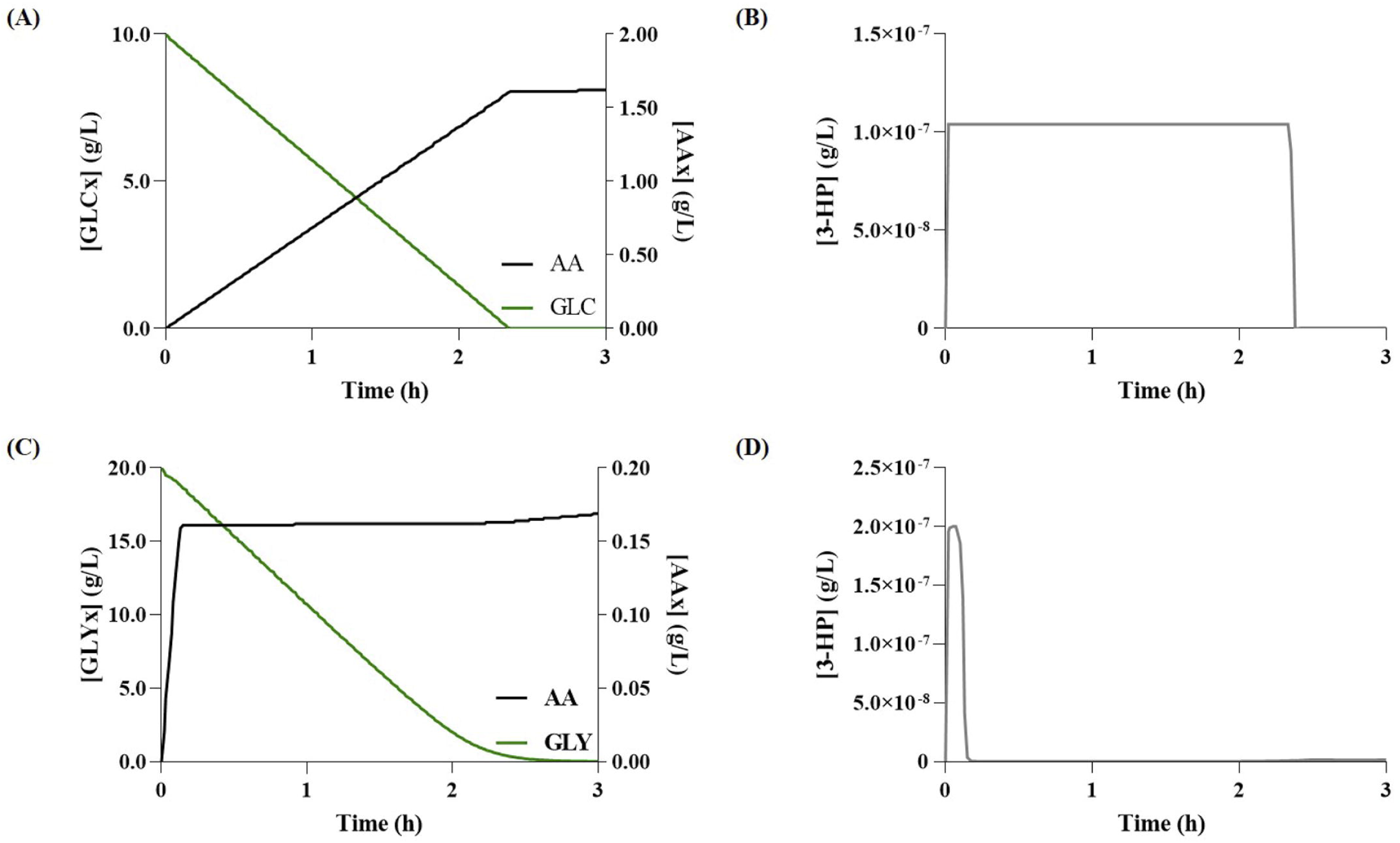
Simulation results for acrylic acid (AA) production via the malonyl-CoA pathway. **(A)** Glucose (GLC) consumption and variation of extracellular AA (AAx) over time; **(B)** Variation of 3-hydroxypropionate (3-HP) concentration over time when using Glucose as carbon source; **(C)** Glycerol (GLYx) consumption and variation of extracellular AAx over time; **(D)** Variation of 3-HP concentration over time when using Glycerol as carbon source.

**Fig 6.**
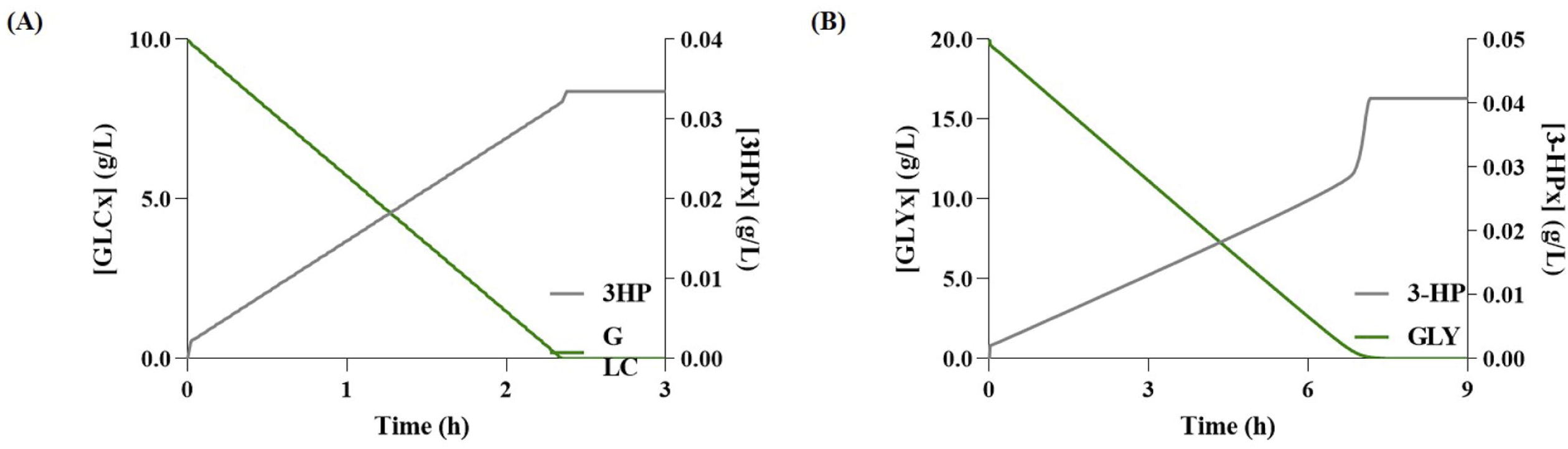
Simulation results for 3-hydroxypropionate (3-HP) production via the *β*-alanine pathway. **(A)** Glucose (GLCx) consumption and variation of extracellular 3-HP (3-HPx) over time; **(B)** Glycerol (GLYx) consumption and variation of 3-HPx over time.

Regarding the Glu-Mcoa models, the 3-HP production predictions are very similar to those found in the literature (Table 4). However, the models failed to predict the production of AA, as Liu and Liu, (2016) [7] reported the accumulation of 3-HP, which was not replicated by the model (Fig 6).

**Table 4.** Summarised results of acrylic acid production for the mutant strains developed for the glycerol, malonyl-CoA, and *β*-alanine models, using 10 g/L of glucose as substrate.

| Pathway | Strain | Predicted<br>Titer (g/L) |
| --- | --- | --- |
| Glycerol | Mutant 0 | 0.16 |
|  | Mutant 1 | 3.11 |
| Malonyl-CoA | Mutant 0 | 1.62 |
|  | Mutant 1 | 3.11 |
| β-alanine | Mutant 0 | 0.027 |
|  | Mutant 1 | 0.97 |

Finally, regarding the *β*-alanine pathway, as shown in Fig 7 and Fig 8, the *β*-alanine model set to use glycerol as carbon source (Gly-Ba) predicted the production of 0.041 g/L of 3-HP and 0.034 g/L of AA. The model set to use glucose as a carbon source (Glu-Ba) predicted the production of 0.033 g/L of 3-HP and 0.027 g/L of AA. Again, the models did not predict the accumulation of 3-HP that, although desirable, is not realistic. Although simulation results indicate a slight advantage towards using glycerol as a carbon source, there seems to be a consensus in literature towards using glucose as carbon source, as to the best of our knowledge, no studies using the glycerol pathway have been reported. When compared to previous pathways, both carbon sources produced a considerably lower concentration of AA in this pathway.

**Fig 7.**
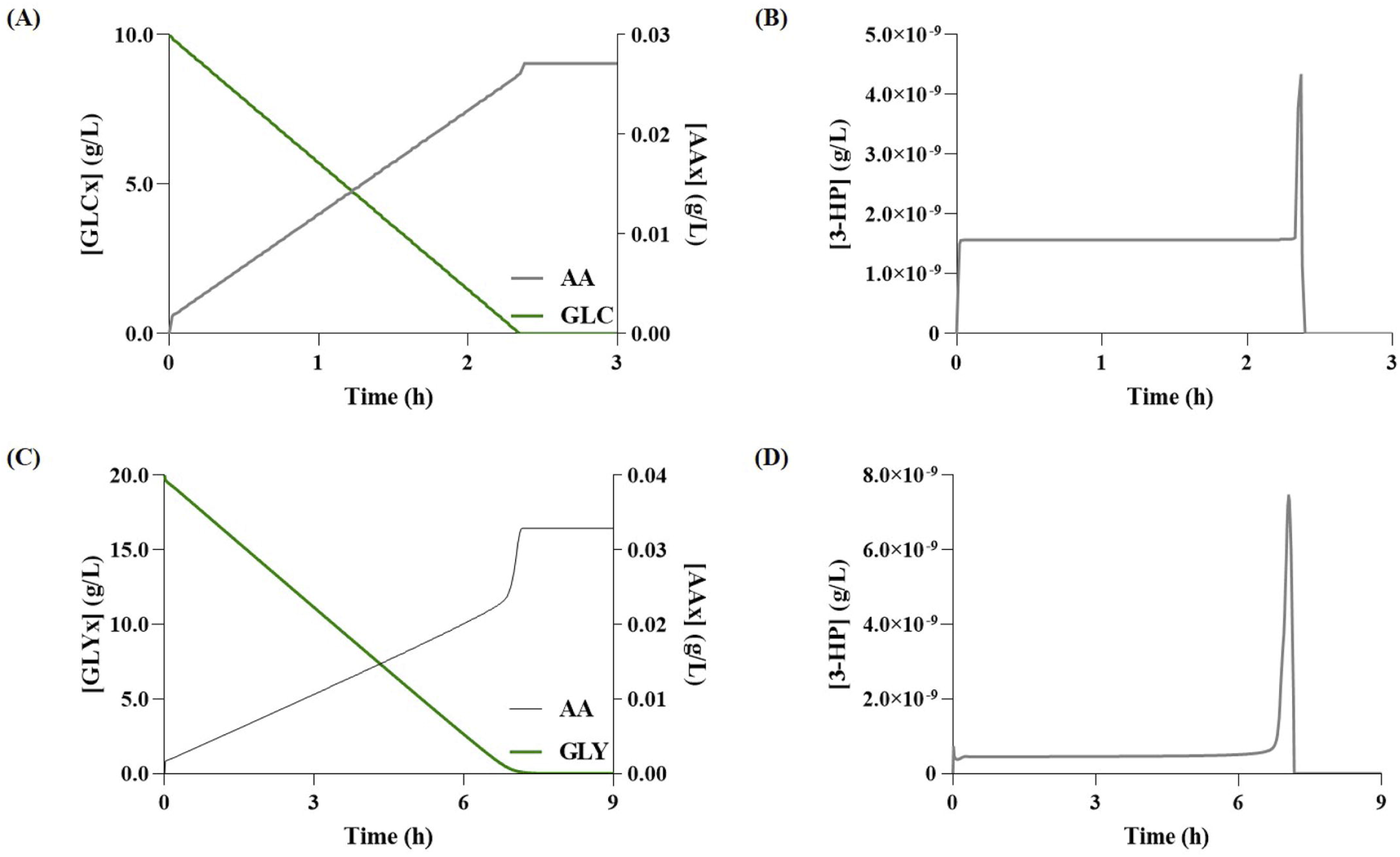
Simulation results for acrylic acid (AA) production via the *β*-alanine pathway. **(A)** Glucose (GLC) consumption and variation of extracellular AA (AAx) over time; **(B)** Variation of 3-hydroxypropionate (3-HP) concentration over time when using Glucose as carbon source; **(C)** Glycerol (GLYx) consumption and variation of extracellular AAx over time; **(D)** - Variation of 3-HP concentration over time when using Glycerol as carbon source.

**Fig 8.**
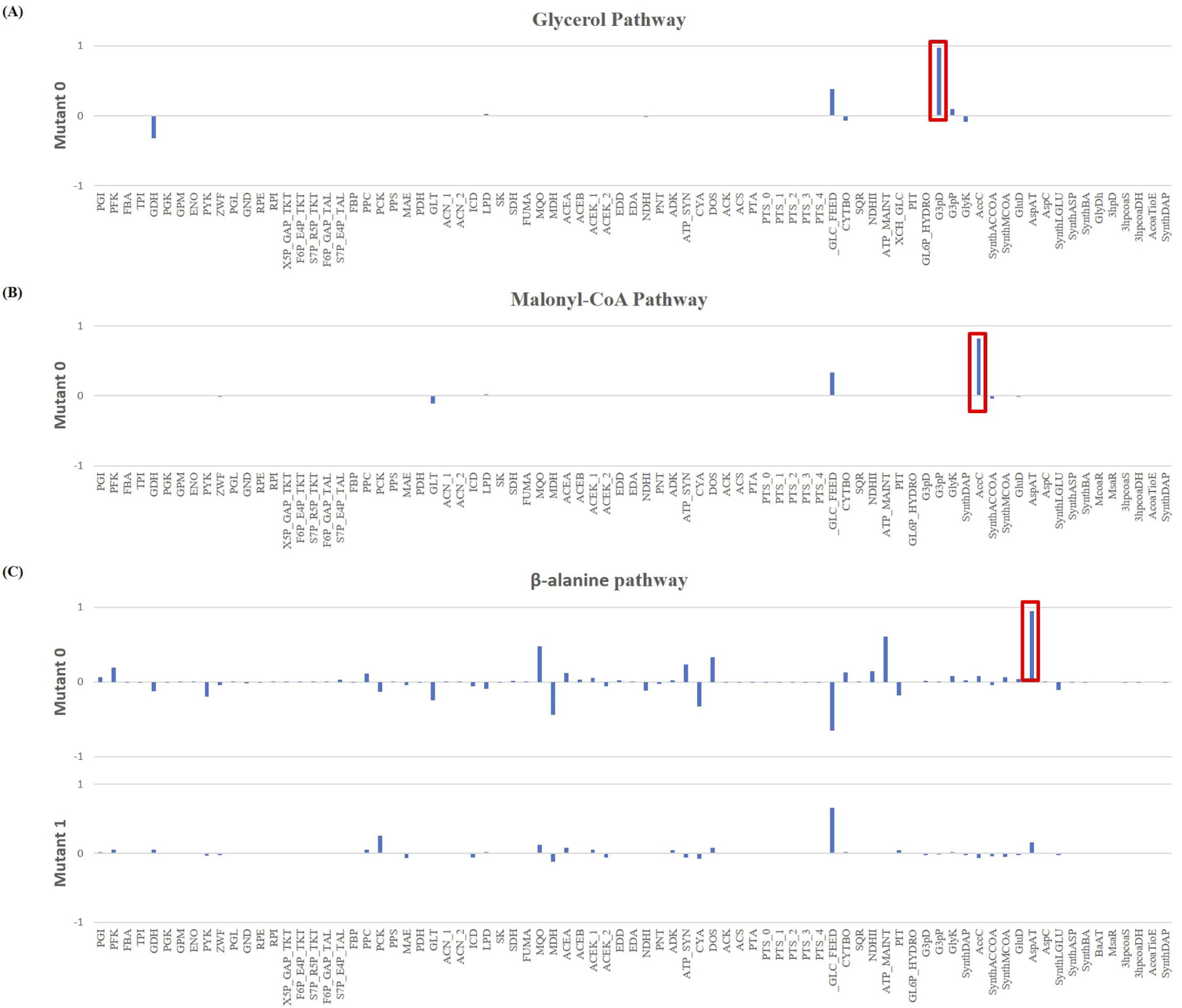
Flux Control Coefficients (FCC) results for acrylic acid formation, where the reaction with the most impact in the yield is highlighted in red. **(A)** Results for the glycerol pathway. According to the coefficients, the reaction with the most impact is the glycerol-3-phosphate dehydrogenase (G3pD), which due to its positive FCC is a potential target for overexpression; **(B)** Results for the malonyl-CoA pathway. The results showed that the acetyl-CoA carboxylase (AccC) is a potential bottleneck in the pathway due to the positive FCC; hence, another target for overexpression; **(C)** Results for the *β*-alanine pathway. The highest FCC was for the aspartate aminotransferase (AspAT) which appears to be an ideal target for an overexpression.

The *β*-alanine pathway is the least studied, with very few reports, which might be associated with the fact that such studies reported significantly lower yields, when compared with the previous pathways [17]. Indeed, only one study was found concerning 3-HP synthesis in *E. coli* using batch cultures [11], while studies in which AA is produced through this route are yet to be published. Despite that, the *β*-alanine model showed promising predictions, at least when comparing to the results published by Song et al. (2016) [11] (Table 5).

**Table 5.**
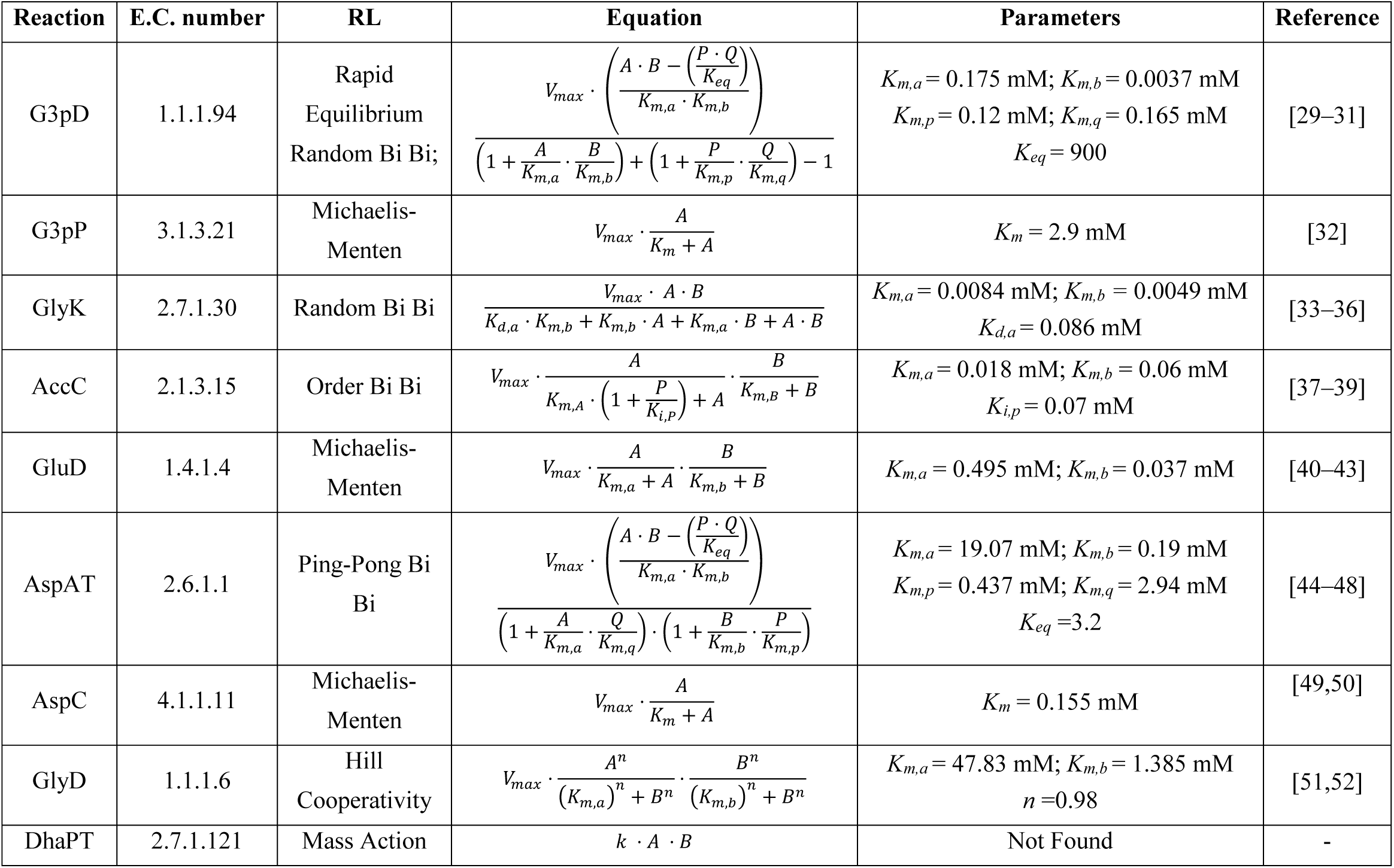
Rate Law (RL) equations, kinetic parameters and the respective references for each reaction that belong to the native metabolism of *Escherichia coli*.

| Reaction | E.C. number | RL | Equation | Parameters | Reference |
| --- | --- | --- | --- | --- | --- |
| G3pD | 1.1.1.94 | Rapid<br>Equilibrium<br>Random Bi Bi; | $V_{max} \cdot \frac{\left( A \cdot B - \left( \frac{P \cdot Q}{K_{eq}} \right) \right)}{\left( 1 + \frac{A}{K_{m,a}} \cdot \frac{B}{K_{m,b}} \right) + \left( 1 + \frac{P}{K_{m,p}} \cdot \frac{Q}{K_{m,q}} \right) - 1}$ | $K_{m,a} = 0.175 \text{ mM}; K_{m,b} = 0.0037 \text{ mM}$<br>$K_{m,p} = 0.12 \text{ mM}; K_{m,q} = 0.165 \text{ mM}$<br>$K_{eq} = 900$ | [29–31] |
| G3pP | 3.1.3.21 | Michaelis-Menten | $V_{max} \cdot \frac{A}{K_m + A}$ | $K_m = 2.9 \text{ mM}$ | [32] |
| GlyK | 2.7.1.30 | Random Bi Bi | $\frac{V_{max} \cdot A \cdot B}{K_{d,a} \cdot K_{m,b} + K_{m,b} \cdot A + K_{m,a} \cdot B + A \cdot B}$ | $K_{m,a} = 0.0084 \text{ mM}; K_{m,b} = 0.0049 \text{ mM}$<br>$K_{d,a} = 0.086 \text{ mM}$ | [33–36] |
| AccC | 2.1.3.15 | Order Bi Bi | $V_{max} \cdot \frac{A}{K_{m,A} \cdot \left( 1 + \frac{P}{K_{i,p}} \right) + A} \cdot \frac{B}{K_{m,B} + B}$ | $K_{m,a} = 0.018 \text{ mM}; K_{m,b} = 0.06 \text{ mM}$<br>$K_{i,p} = 0.07 \text{ mM}$ | [37–39] |
| GluD | 1.4.1.4 | Michaelis-Menten | $V_{max} \cdot \frac{A}{K_{m,a} + A} \cdot \frac{B}{K_{m,b} + B}$ | $K_{m,a} = 0.495 \text{ mM}; K_{m,b} = 0.037 \text{ mM}$ | [40–43] |
| AspAT | 2.6.1.1 | Ping-Pong Bi<br>Bi | $V_{max} \cdot \frac{\left( A \cdot B - \left( \frac{P \cdot Q}{K_{eq}} \right) \right)}{\left( 1 + \frac{A}{K_{m,a}} \cdot \frac{Q}{K_{m,q}} \right) \cdot \left( 1 + \frac{B}{K_{m,b}} \cdot \frac{P}{K_{m,p}} \right)}$ | $K_{m,a} = 19.07 \text{ mM}; K_{m,b} = 0.19 \text{ mM}$<br>$K_{m,p} = 0.437 \text{ mM}; K_{m,q} = 2.94 \text{ mM}$<br>$K_{eq} = 3.2$ | [44–48] |
| AspC | 4.1.1.11 | Michaelis-Menten | $V_{max} \cdot \frac{A}{K_m + A}$ | $K_m = 0.155 \text{ mM}$ | [49,50] |
| GlyD | 1.1.1.6 | Hill<br>Cooperativity | $V_{max} \cdot \frac{A^n}{(K_{m,a})^n + B^n} \cdot \frac{B^n}{(K_{m,b})^n + B^n}$ | $K_{m,a} = 47.83 \text{ mM}; K_{m,b} = 1.385 \text{ mM}$<br>$n = 0.98$ | [51,52] |
| DhaPT | 2.7.1.121 | Mass Action | $k \cdot A \cdot B$ | Not Found | - |

### Optimisation Strategies

Ideally, all models capable of producing AA should have been optimised. However, as mentioned before, the Gly-Mcoa model presented issues with dihydroxyacetone accumulation, not being further used in this work. Additionally, the Gly-Gly and Gly-Ba models proved to be unstable when performing the metabolic control analysis (MCA), preventing the flux control coefficient (FCC) ascertainment, due to the lack of a steady-state. Henceforth, only the models designed to use glucose as a carbon source (Glu-Gly, Glu-Mcoa, Glu-Ba) were optimised.

Starting with the glycerol model, the MCA showed that the enzyme with the highest FCC, and thus a more significant influence on AA production, was the G3pD (Fig 9A), which is responsible for converting dihydroxyacetone phosphate into glycerol-3-phosphate. The reaction catalysed by this enzyme is a potential bottleneck in the pathway, thus a target for overexpression. Using COPASI’s optimisation task, Mutant Glu-Gly 1 was created. This mutant included an *in silico* overexpression of the selected enzyme, in which the *V*_*max*_ of the enzyme was set to 1.392 mM/s, representing a nearly 45-fold increase that resulted in the production of 3.11 g/L of AA (Fig 10A). A subsequent MCA was performed on Mutant Glu-Gly 1, aiming at further optimising the AA production yields. However, the model could not reach a steady-state; hence, the FCCs were not available, thus terminating the optimisation of this model.

**Fig 9.**
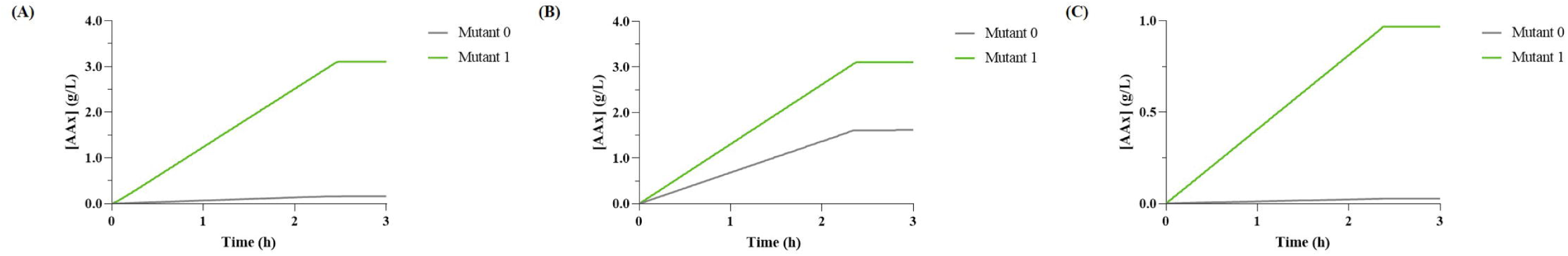
Comparison between the original acrylic acid (AA) production with the results obtained for the mutants developed with the optimisation strategies identified. **(A)** AA production using glycerol pathway. Mutant 0 represents the model with the heterologous pathway, and Mutant 1 the same model with a 45-fold increase in the *V*_*max*_ of the glycerol-3-phosphate dehydrogenase (G3pD) reaction; **(B)** AA production using malonyl-CoA pathway. Mutant 0 represents the model with the heterologous pathway, and Mutant 1 the same model with a 2.5-fold increase in the *V*_*max*_ of the acetyl-CoA carboxylase (AccC); **(C)** AA production for the *β*-alanine pathway. Mutant 0 represents the model with the heterologous pathway, and Mutant 1 the same model with a 50-fold increase in the *V*_*max*_ of the aspartate aminotransferase (AspAT).

**Fig 10.**
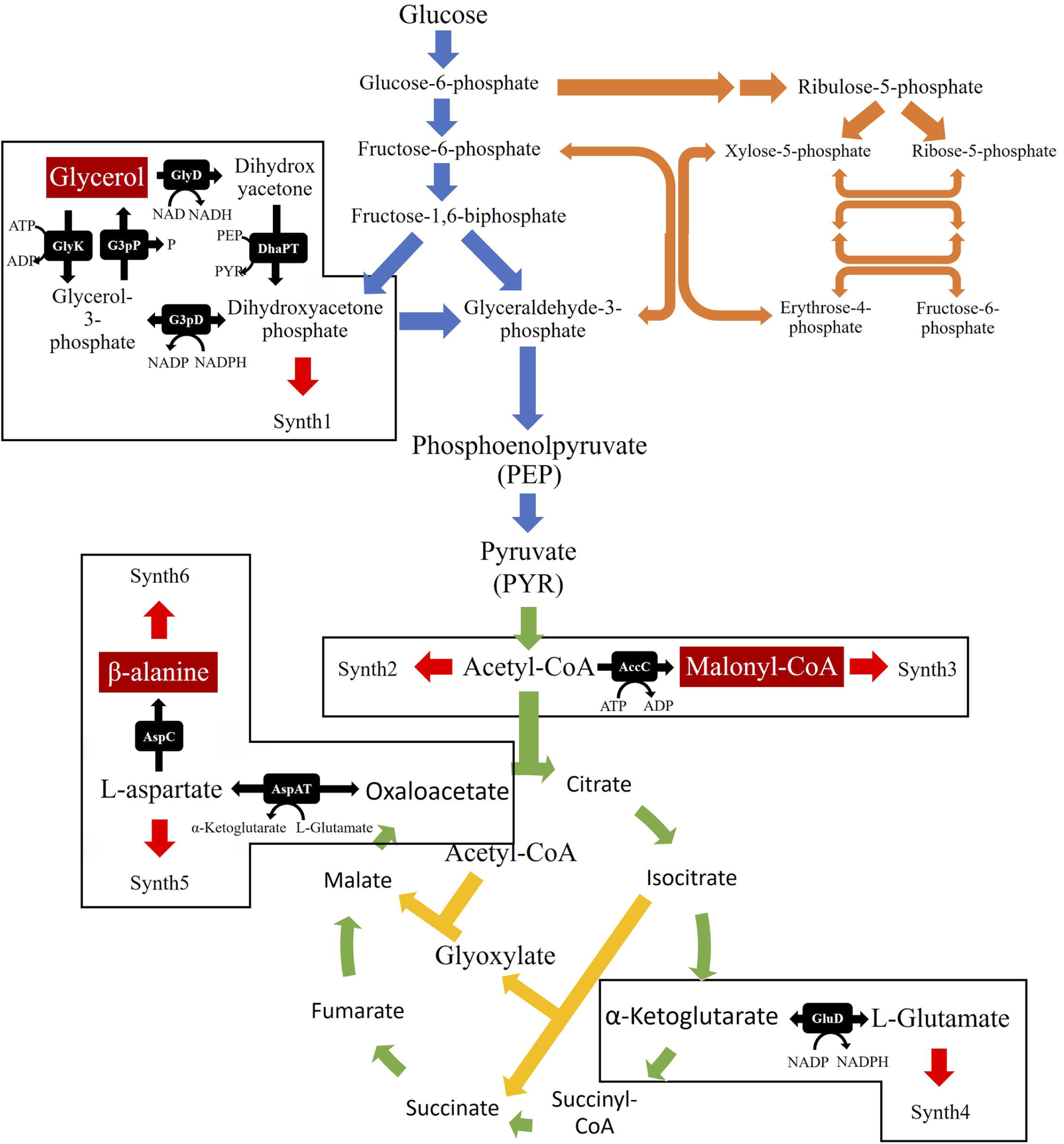
Representation of the central carbon metabolism of *Escherichia coli* and the reactions added to the kinetic model. The reactions depicted by the blue, orange, green and yellow arrows represent, respectively, the glycolysis, pentose-phosphate pathway, tricarboxylic acid cycle and the glyoxylate shunt, which are all present in the original model. The black arrows represent the nine reactions that were added to the model. Finally, red arrows depict the Synth reactions added to account for the presence of the newly added metabolites in other pathways

In the malonyl-CoA pathway model, the FCCs identified one potential overexpression target, the AccC (Fig 9B), which is responsible for the conversion of acetyl-CoA to malonyl-CoA. The optimum *V*_*max*_ for this enzyme was found to be 0.568 mM/s, corresponding to approximately a 2-fold overexpression. Mutant Glu-Mcoa 1 was able to produce a 3.11 g/L of AA, which corresponds to a 1.5-fold higher yield than Mutant Glu-Mcoa 0 (Fig 10B). Unfortunately, once again, a second iteration revealed that the model was unable to reach a steady-state. Thus, it was not possible to identify other potential targets using this methodology.

Regarding the *β*-alanine pathway, the MCA showed that the model has several reactions affecting the AA yield. However, the reaction with the most significant impact is catalysed by AspAT enzyme (Fig 9C). This reaction allows converting oxaloacetate and *L-*glutamate into aspartate, which is in turn converted to *β*-alanine. This reaction has a positive coefficient, which indicates that it is a potential bottleneck impairing the downstream flux towards the heterologous pathway; thus, the optimisation goal was to overexpress this enzyme. COPASI estimated that a 50-fold overexpression was ideal for maximising the production yields, resulting in a *V*_*max*_ of 127.4869 mM/s. With this change, the predicted AA production was 0.97 g/L, which corresponds to a 28-fold increase (Fig 10C). A subsequent MCA revealed that the main limiting factor to AA production in Mutant Glu-Ba 1 was the amount of glucose provided to the model; hence the optimisation was terminated with only one target identified. Similarly, the AspC gene, which is responsible for the production of *β*-alanine, can also be considered as a limiting factor for pathway flux, thus becoming a target for optimisation, as the *V*_*max*_ was increased for the *β*-alanine model to work correctly (S1 Appendix, section 1.1).

The goal of these optimisations was to provide guidelines that can be later implemented *in vivo* and not predict the AA production accurately. Nonetheless, the final concentrations that were obtained with the new mutants were used to compare with previous results. As shown in Table 6, the same concentration of AA (3.11 g/L) was produced by the glycerol and malonyl-CoA pathways. The *β*-alanine pathway also showed a substantial yield increase (0.97 g/L). However, the value is still considerably lower than the obtained with remaining pathways.

**Table 6.** Rate Law (RL) equations, kinetic parameters and the respective references for each reaction of the three heterologous pathways (glycerol, malonyl-CoA and *β*-alanine) required to produce acrylic acid.

| Reaction | E.C. number | RL | Equation | Parameters | Reference |
| --- | --- | --- | --- | --- | --- |
| GlyDH | 4.2.1.28 | Specific Activation | $\frac{E \cdot K_{cat} \cdot A \cdot Activator}{K_{m,a} \cdot K_a + (K_{m,a} + A) \cdot Activator}$ | $K_{cat} = 0.0621 \text{ s}^{-1}$; $K_m = 6.15 \text{ mM}$<br>$K_a = 0.008 \text{ mM}$ | [53,54] |
| 3hpaD | 1.2.1.99 | Michaelis-Menten | $\frac{E \cdot K_{cat} \cdot A \cdot B}{K_{m,a} \cdot K_{m,b} + K_{m,b} \cdot A + K_{m,a} \cdot B + A \cdot B}$ | $K_{cat} = 16.73 \text{ s}^{-1}$; $K_{m,a} = 0.39 \text{ mM}$<br>$K_{m,b} = 1.3 \text{ mM}$ | [55] |
| McoaR | 1.2.1.75 | Michaelis-Menten | $\frac{E \cdot K_{cat} \cdot A \cdot B}{K_{m,a} \cdot K_{m,b} + K_{m,b} \cdot A + K_{m,a} \cdot B + A \cdot B}$ | $K_{cat} = 50 \text{ s}^{-1}$; $K_{m,a} = 0.3 \text{ mM}$<br>$K_{m,b} = 0.03 \text{ mM}$ | [56,57] |
| MsaR | 2.6.1.19 | Michaelis-Menten | $\frac{E \cdot K_{cat} \cdot A \cdot B}{K_{m,a} \cdot K_{m,b} + K_{m,b} \cdot A + K_{m,a} \cdot B + A \cdot B}$ | $K_{cat} = 115 \text{ s}^{-1}$; $K_{m,a} = 0.07 \text{ mM}$<br>$K_{m,b} = 0.07 \text{ mM}$ | [58] |
| BaTA | 1.1.1.298 | Ping-Pong Bi Bi | $\frac{E \cdot K_{cat} \cdot A \cdot B}{K_{m,b} \cdot A + K_{m,a} \cdot B \cdot \left(1 + \frac{B}{K_{i,B}}\right) + A \cdot B}$ | $K_{cat} = 47.4 \text{ s}^{-1}$; $K_{m,a} = 5.8 \text{ mM}$<br>$K_{m,b} = 1.07 \text{ mM}$; $K_{i,b} = 10.2 \text{ mM}$ | [59] |
| 3hpcoaS | 6.2.1.36 | Michaelis-Menten | $E \cdot K_{cat} \cdot \frac{A}{K_{m,a} + A} \cdot \frac{B}{K_{m,b} + B} \cdot \frac{C}{K_{m,c} + C}$ | $K_{cat} = 36 \text{ s}^{-1}$; $K_{m,a} = 0.015 \text{ mM}$<br>$K_{m,b} = 0.01 \text{ mM}$; $K_{m,c} = 0.05 \text{ mM}$ | [60] |
| 3hpcoaDH | 4.2.1.116 | Michaelis-Menten | $\frac{E \cdot K_{cat} \cdot A}{K_m + A}$ | $K_{cat} = 96 \text{ s}^{-1}$; $K_m = 0.06 \text{ mM}$ | [61] |
| AcoaTioE | 3.1.2.20 | Michaelis-Menten | $\frac{E \cdot K_{cat} \cdot A}{K_m + A}$ | $K_{cat} = 0.55 \text{ s}^{-1}$; $K_m = 0.167 \text{ mM}$ | [62] |

The enzyme concentration ( $E$ ) value used for all the reactions was 100 mM.

### Conclusion

In conclusion, these models seem to be more accurate in predicting 3-HP synthesis because the *V*_*max*_ for the heterologous enzymes has calculated in excess so that it would not limit the reaction flux. Moreover, the models also exhibited issues associated with the assimilation of glycerol and *β*-alanine production. An effort was put forward towards finding proteomics data that included the absolute quantification of such enzymes to solve these problems. However, no quantification data was found; therefore, it will be important in the future to seek such data or, in the lack of new data, to determine it experimentally. Additionally, an overall best carbon source does not emerge from this work. Instead, the answer is specific to the selected pathway. When comparing the three bio-based routes, the results suggest that the glycerol pathway leads to the highest yields, when combined with the used of glycerol as a carbon source. However, a relevant caveat must be recalled. This pathway involves a reaction that relies on the presence of vitamin B_12_, which represents a significant economic disadvantage at an industrial-scale production [8,9]. Hence, to make this route economically viable, either the yield must be significantly improved to overcome the cost of the vitamin supplementation, or a cheaper way to produce B_12_ must be found. Therefore, despite producing less AA, it seems to be beneficial to use the malonyl-CoA route, as it provided the second-highest yield, does not need any vitamin supplementation, and there is still room for optimisation [16–18]. Finally, the model optimisation revealed four targets for overexpression with the goal of increasing the bioavailability of the intermediaries (glycerol, malonyl-CoA, or *β*-alanine). Nevertheless, these models only comprise the CCM. Thus, other strategies to force additional flux towards the heterologous pathway may prove effective. Finally, after the model optimisation, the final step should be the validation with *in vivo* experiments to further confirm the results herein gathered.

## Materials and Methods

### Kinetic Modelling

The dynamic model developed by Millard et al. (2017) [19] was extended to include the missing pathways associated with 3-HP and AA production. This process was divided into three steps, the extension of the CCM to include the production of the three intermediary compounds (glycerol, malonyl-CoA, and *β*-alanine), the subsequent production of 3-HP, and the production of AA.

### Parameter Selection

Kinetic equations and their respective parameters were retrieved from the available literature. Databases like BioCyc [20], BRENDA [21], Sabio-RK [22] and eQuilibrator [23] were used to identify the kinetic mechanism of each enzyme and obtain their respective parameters. Furthermore, instead of a single value, the average of all parameters found for each enzyme, excluding outliers, was used to obtain a better representation. A summary of the values considered when calculating the mean value for each reaction is presented in S2 Appendix.

Regarding the parameters required to describe a reaction, the maximal rate (*V*_*max*_) is usually not reported in the literature. Unfortunately, this parameter is highly dependent on the specificity of the assay conditions. Hence, the specific activity or the turnover (a.k.a. *K*_*cat*_) are reported instead. Two distinct methods were used, as a workaround, to estimate values for this parameter. Method 1, adapted from the work of Chassagnole and colleagues [24], was used for reactions belonging to *E. coli*’s native metabolism. Initially, a steady-state flux distribution is determined for the original kinetic model. Then, a genome-scale model of *E. coli* K-12 MG1655 (in this case iML1515) [25] is used to predict the flux of the new reactions. For this purpose, the common reactions between the kinetic model and the stoichiometric model are constrained to the previously determined flux distribution (±0.01 mM/s). Then, a flux variability analysis [26] is performed to estimate the maximum flux (*v*). By equalising *v* to the respective rate law (*V*_*max*_ × *F (X,K)*), the following equation is obtained:

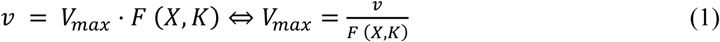

in which *X* is a vector of parameters, and *K* a vector of steady-state concentrations for the metabolites involved in the respective reaction. Furthermore, notice that for newly added metabolites, the steady-state concentration was assumed to be 1 mM. An example of these calculations and the resulting *V*_*max*_ values are presented in the S1 Appendix, section 1.2.

Method 2 was used for reactions of the heterologous pathways. In this method, the *V*_*max*_ was estimated assuming that the total concentration of enzyme was in surplus (100 mM), thus calculating this parameter as shown in equation 2:

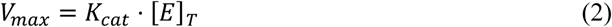

### Extension of the Central Carbon Metabolism (CCM)

The production of glycerol, malonyl-CoA, and *β*-alanine had to be included in the model to insert the three heterologous pathways (Fig 2). The reactions catalysed by the glycerol-3-phosphate dehydrogenase (G3pD) and glycerol-3-phosphate phosphatase (G3pP) enzymes are required to produce glycerol from dihydroxyacetone phosphate. Considering that this route has to be reversible to use glycerol as a carbon source, reactions catalysed by the glycerol kinase (GlyK), glycerol dehydrogenase (GlyD) and the dihydroxyacetone phosphate transferase (DhaPT) enzymes were included too (Fig 2). As shown in Fig 2, one reaction is required to obtain malonyl-CoA, namely the reaction catalysed by the acetyl-CoA carboxylase (AccC) enzyme. Finally, three reactions were included for the *β*-alanine pathway. Two of these, promoted by the aspartate aminotransferase (AspAT) and the aspartate carboxylase (AspC) enzymes, are required for the production of *β*-alanine. A reaction, catalysed by the *L-*glutamate dehydrogenase (GluD) enzyme, is used to produce glutamate, which is required by the AspAT to produce aspartate (Fig 2) [20,27].

All reactions and respective stoichiometry are shown below:

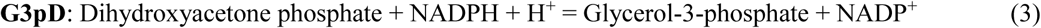

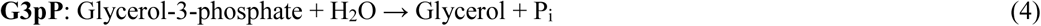

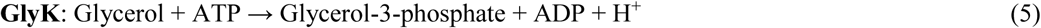

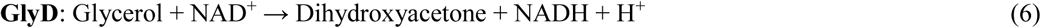

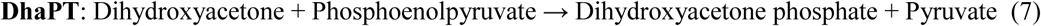

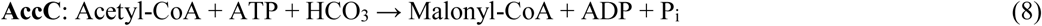

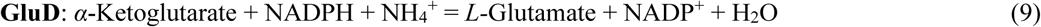

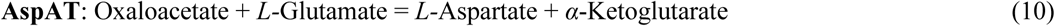

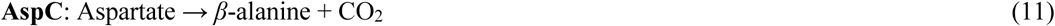

Furthermore, the kinetic law equation and the respective parameters for each of the previously described reactions are presented in Table 1.

An additional set of pseudo-reactions was included in the model; the Synth reactions (Fig 2). These reactions, inspired by the work of Chassagnole et al. (2002) [24] and Machado et al. (2014) [28], are used to represent the pathways involved in the breakdown of the newly added metabolites. Mass action kinetics was assumed for these reactions and, using the same principle as Method 1, the sum of all fluxes from the reactions that metabolise each metabolite in the stoichiometric model was used to determine the *k* values for each synth reaction. The resulting *k* values are available in the S1 Appendix, section 1.2.

### Pathways for Acrylic Acid Production

The following step was to insert the three heterologous pathways to produce AA separately into the extended CCM model. All pathways encompass two different phases, the production of an intermediary compound, namely 3-HP, and subsequent production of AA (Fig 1).

The first phase involves two different enzymes in each pathway. Regarding the glycerol pathway, such enzymes are the glycerol dehydratase (GlyDH) and the 3-hydroxypropionaldehyde dehydrogenase (3hpaD). In the malonyl-CoA pathway, the malonyl-CoA reductase (McoaR) enzyme is responsible for the production of malonic semialdehyde (MSA), which is then converted into 3-HP by the malonic semialdehyde reductase (MsaR). Regarding the β-alanine pathway, the *β*-alanine aminotransferase (BaAT) enzyme promotes the conversion of β-alanine together with *α*-ketoglutarate into *L*-glutamate and MSA. The later is then converted into 3-HP by the MsaR.

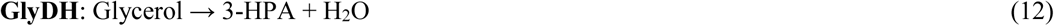

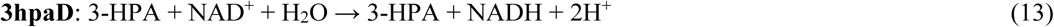

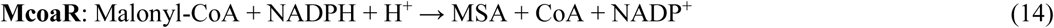

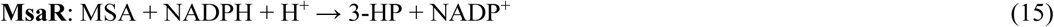

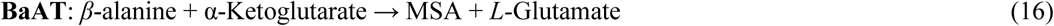

The final step is to convert the newly formed 3-HP into AA. This process involves the production of 3-hydroxypropionyl-CoA (3-HP-CoA) by the 3-hydroxypropionyl-CoA synthase (3hpcoaS), the subsequent formation of acrylyl-CoA (AA-CoA) by the 3-hydroxypropionyl-CoA dehydratase (3hpcoaDH), and finally, the production of AA by the acrylyl-CoA thioesterase (AcoaTioE) enzyme, as shown in Fig 1. The stoichiometry of these reactions is likewise shown below, and the respective kinetic parameters presented in Table 2.

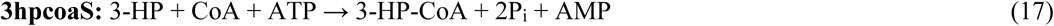

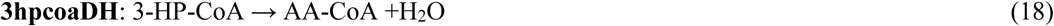

The enzyme concentration (*E*) value used for all the reactions was 100 mM.

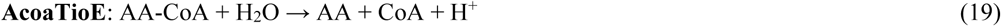

Four models were created for each of the three pathways resulting in a total of twelve distinct models. To be more precise, for each pathway, models to produce 3-HP or AA, from glucose or glycerol as carbon sources, were put forward. All models can be found at https://cutt.ly/aaKineticModels.

### Time Course Simulation

Time course simulations were performed to assess 3-HP and AA production over time, using the deterministic method (LSODA) from COPASI [63], with a duration of three or six hours, to allow the consumption of all available carbon source. Since the available carbon sources have a different number of carbons, the initial concentration of such molecules had to ensure that the amount of carbon provided to the model was the same. Thus, the initial concentrations for glucose and glycerol were 55.5 mM (10 g/L) and 217.2 mM (20 g/L), respectively, which allowed comparing the three pathways for each carbon source. The model was assessed to available literature regarding these pathways, in which the simulations’ initial concentration of the carbon source was set to replicate the initial conditions of published results.

### Optimisation Strategies

The first step was to determine the flux control coefficients (FCC), through a metabolic control analysis (MCA). Initially, a feed and drains for glycerol, malonyl-CoA, *β*-alanine, and AA were included to replicate a continuous model and find a valid steady-state, which is required to determine the FCCs. These coefficients reflect the level of control that each reaction has over the formation of AA. Nevertheless, a batch version of the model was used for the optimisation. The optimisation was performed using the automated optimisation tool provided by COPASI [63]. Here, the algorithm proposed new *in silico* mutant strains, in which the reaction with most influence was either over-expressed, under-expressed, or knocked-out through a change in the *V*_*max*_, according to the respective coefficient. The goal of the optimisation was not to meticulously predict the final concentration of AA, but rather to find promising targets for optimisation. Hence, the changes in the *V*_*max*_ were limited to 50 times the original value to allow overcoming the influence of such reaction *in silico*, while not impairing the *in vivo* implementation.

The objective function was the maximisation of AA concentration in the time course task. After creating new mutants, the process was repeated to optimise the mutant strains further. The task eventually stopped when either the glucose feed was limiting the production of AA, the limiting reaction was already optimised, or the system could no longer reach a stable steady-state point during the MCA.

## Acknowledgments

This study was supported by the Portuguese Foundation for Science and Technology (FCT) under the scope of the strategic funding of UIDB/04469/2020 unit and BioTecNorte operation (NORTE-01-0145-FEDER-000004) funded by the European Regional Development Fund under the scope of Norte2020 - Programa Operacional Regional do Norte. This article is also a result of the project 22231/01/SAICT/2016: “Biodata.pt – Infraestrutura Portuguesa de Dados Biológicos”, supported by Lisboa Portugal Regional Operational Programme (Lisboa2020), under the PORTUGAL 2020 Partnership Agreement, through the European Regional Development Fund (ERDF).

## Author Contributions

AO performed the *in silico* simulations, analysed the results and draft the manuscript; EF and LR provided feedback and suggestions on the manuscript; JR and OD assisted in the study conception and design, and provided feedback and suggestions on the manuscript. All authors have read and agreed to the published version of the manuscript.

## Supporting information

**S1 Appendix. Supplementary results and parameter adjustments.** Explanations behind parameter adjustments adopted to circumvent simulation issues, and results for the Vmax calculation using method 1 and method 2.

**S2 Appendix. Kinetic parameters.** This file will present all the kinetic parameters and equations used to model AA production.

## Notes

https://cutt.ly/aaKineticModels

### Competing Interest Statement

The authors have declared no competing interest.

### Summary of Updates

The changes were mainly made in the structure and template of the manuscript, as it was adapted to fit the submission requirements. Some changes include changing the order of the sections, adding line count to the document, changing the font and font size, among others. Also, the images were placed at the end of the PDF file, as well as the supplementary information. Regarding the results for the simulation of the β-alanine pathway, we found that the values for acrylic acid production were being presented as results of 3-hydroxypropionate and vice-versa, which had to be corrected. Finally, several typos that were found during this process were also fixed.

