## Supporting information for "A kinetic model of the central carbon metabolism for acrylic acid production in *Escherichia coli*"

### S1 Appendix

#### 1 Supplementary Results

##### 1.1 Simulation Problems

During the simulations, several issues arised. The first one was related to the $V_{max}$ of the aspartate carboxylase (AspC) enzyme during the CCM extension. The second problem was related by the assimilation of glycerol to the CCM in the glycerol and malonyl-CoA models when using glycerol as carbon source, from which it was only possible to circumvent the first one. In this section these problems will be further explained as well as the methods used to resolve them.

###### 1.1.1 Extension of the Central Carbon Metabolism (CCM)

Even though the simulation of glycerol and malonyl-CoA production was successful, the production of $\beta$ -alanine exhibited a major setback. The extremely low $V_{max}$ of the AspC enzyme (S2 Table 1) was limiting the amount of $\beta$-alanine produced (S2 Fig 1A). The $V_{max}$ was calculated with the flux obtained from the stoichiometric model simulation (Method 1), which resulted in a relevant bottleneck. Thus, this parameter was recalculated using the method developed for the heterologous pathway reactions parameters (Method 2), shifting the $\beta$-alanine flux production limits towards L-aspartate formation. According to Ramjee et al. (1997) [49], the $K_{cat}$ value for this enzyme was 0.57 $s^{-1}$ [118]. Therefore, the $V_{max}$ value was set to 57 mM/s in all the models, which drastically altered the production of $\beta$-alanine (S2 Fig 1B).

**S1 Fig 1. Variation of *β*-alanine (BA) production over time.**

(A) *β*-alanine concentration using the $V_{max}$ for the aspartate carboxylase (AspC) enzyme calculated using Method 1 ($1.15x10^{-05}$ mM/s). (B) *β*-alanine concentration using the $V_{max}$ for the AspC calculated using Method 2 (57 mM/s).

##### 1.1.2 Glycerol Model

Issues were found in the models created for modeling 3-HP and AA production, when glycerol was used as carbon source. The carbon flow toward the CCM stopped seconds after the simulation started, leading to the accumulation of 3-HP (S2 Fig 2). After analyzing the system, it was determined that two factors were responsible for this behavior. The first factor is associated with the heterologous pathway, as the $V_{max}$ of all reactions in this pathway was set as not to limit flux through these reactions. Therefore, these reactions would uptake most of the available glycerol, thus limiting the amount of carbon towards the CCM, which eventually led to the depletion of crucial cofactors for AA synthesis, such as $NAD^+$ and ATP. The second factor is associated with the $NAD^+$ affinity to enzymes glycerol dehydrogenase (GlyD) and 3-hydroxypropionaldehyde dehydrogenase (3hpaD). The $K_m$ value in the GlyD is over tenfold higher than in 3hpaD, thus the low concentration of $NAD^+$ available was mainly used by the 3hpaD, which blocked the flux towards the CCM, exacerbating the energy production problem even further.

**S1 Fig 2. Results of the time course simulations in the original glycerol model.**

(A) Glycerol (GLY) consumption. (B) Production of 3-hydroxypropionate (3-HP). (C) Acrylic acid (AA) production. (D) Flux of the glycerol dehydrogenase (GlyD).

A couple of hypotheses were devised to circumvent these problems. Because the $NAD^+$ affinity issue was blocking the flux of carbon towards the CCM, the first hypothesis was increasing the affinity of the GlyD enzyme towards $NAD^+$. When comparing the affinity with other enzymes present in the model, this enzyme had a significantly higher value. Therefore, other works that characterized this enzyme were searched, to find an alternative for this $K_m$ value. In the work of Zhang et al. (2010) [52] this enzyme, using another substrate, was described with an affinity towards $NAD^+$ of 0.0165 mM. Thus, the value was updated and simulated in a period of three hours. However, the problem persisted as most glycerol was still going towards the formation of 3-HP (S2 Fig 3).

**S1 Fig 3. Results of the time course simulations in the original glycerol model after the affinity towards $NAD^+$ of the GlyD was changed to 0.0165 mM.**

(A) Glycerol (GLY) consumption. (B) Production of 3-hydroxypropionate (3-HP). (C) Acrylic acid (AA) production. (D) Flux of the glycerol dehydrogenase (GlyD).

The next strategy was increasing the maximal rate of the GlyD to increase the flux in the CCM, allowing the continuous production of AA. Hence, the $V_{max}$ was recalculated according to Method 2, resulting in a value of 4298.4 mM/s. However, even then the majority of glycerol provided was directed towards the heterologous pathway (S2 Fig 4B), and the small amount that was directed towards the CCM was accumulated as dihydroxyacetone because the activity of the dihydroxyacetone phosphate transferase (DhaPT) was not enough to consume that metabolite (S2 Fig 4C).

**S1 Fig 4. Results of the time course simulations in the original glycerol model after the $V_{max}$ of the reaction GlyD was changed to 4298.4 mM/s.**

(A) Glycerol (GLY) consumption. (B) Production of 3-hydroxypropionate (3-HP). (C) Variation of dihydroxyacetone (DHA) concentration. (D) Acrylic acid (AA) production.

The third hypothesis involved decreasing the affinity of the 3hpaD towards $NAD^+$, to improve the activity of the GlyD, and the maximum rate of the glycerol dehydratase (GlyDH) to control the flux that is directed towards to heterologous route. Since no further $K_m$ value for the 3hpaD was found in the literature, it was assumed that this enzyme had a similar affinity as the GlyD ($K_m$ = 0.8 mM). Furthermore, the $V_{max}$ of the GlyDH was set to 0.621 mM/s, representing an enzyme concentration of 10 mM instead of the 100 mM used in Method 2. These changes allowed directing the flux towards the CCM, which resulted in the production of AA without the excessive accumulation of any intermediaries (Fig 3).

#### 1.2 $V_{max}$ Calculation

The $V_{max}$ values for the enzymes that belong to the native metabolism of *Escherichia coli*, calculated using method 1, are presented in S2 Table 1:

**S2 Table 1.** $V_{max}$ values calculated for the reactions required for the extension of the central carbon metabolism.

| **Reaction ID** | **$V_{max}$ (mM/s)** |
|---|---|
| G3pD | 0.0309 |
| G3pP | 0.7741 |
| GlyK | 0.2052 |
| AccC | 0.2914 |
| GluD | 0.1716 |
| AspAT | 2.5548 |
| AspC | $1.15 \times 10^{-05}$ |
| GlyD | 14.7187 |
| DhaPT | 0.0586 |

Using the same principle, the *k* values for the synth reactions were also calculated using the restrained stoichiometric model. The calculated values are presented in S2 Table 2.

**S2 Table 2.** Synth reactions added to the model and respective parameters. These reactions were created for dihydroxyacetone phosphate (DAP), acetyl-CoA (ACCOA), malonyl-CoA (MCOA), L-glutamate (LGLU), L-aspartate (ASP) and *β*-alanine (BA).

| Reaction ID | *k* value (1/s) |
|---|---|
| SynthDAP | 0.02006 |
| SynthACCOA | 0.69799 |
| SynthMCOA | 0.09250 |
| SynthLGLU | 0.38260 |
| SynthASP | 0.05180 |
| SynthBA | 0.00001 |

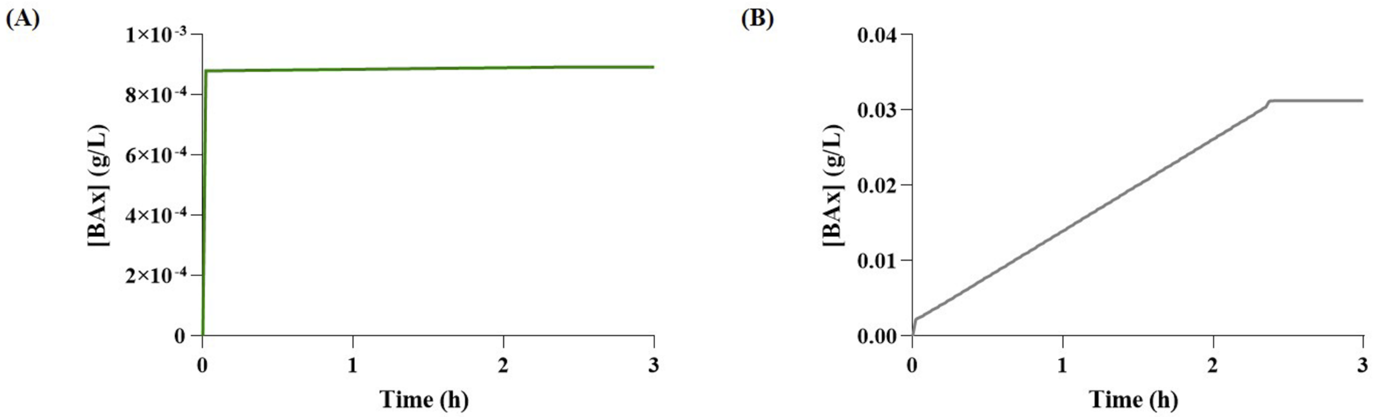
(A)
1×10⁻³
8×10⁻⁴
6×10⁻⁴
4×10⁻⁴
2×10⁻⁴
0
[BAx] (g/L)
0
1
2
3
Time (h)
(B)
0.04
0.03
0.02
0.01
0.00
[BAx] (g/L)
0
1
2
3
Time (h)

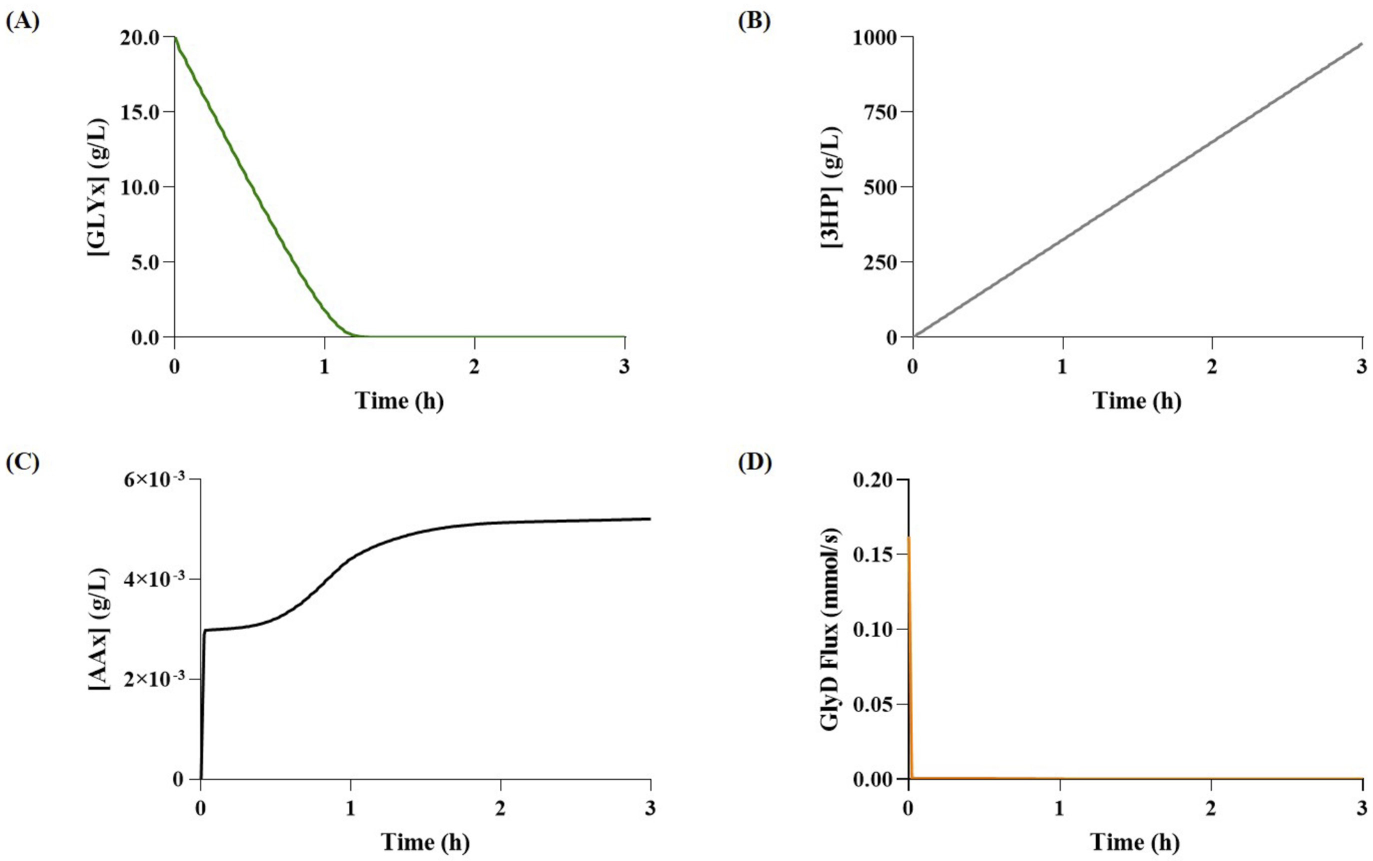

(A)
[GLYx] (g/L)
20.0
15.0
10.0
5.0
0.0
0
1
2
3
Time (h)
(B)
[3HP] (g/L)
1000
750
500
250
0
0
1
2
3
Time (h)
(C)
[AAx] (g/L)
6×10^-3
4×10^-3
2×10^-3
0
0
1
2
3
Time (h)
(D)
GlyD Flux (mmol/s)
0.20
0.15
0.10
0.05
0.00
0
1
2
3
Time (h)

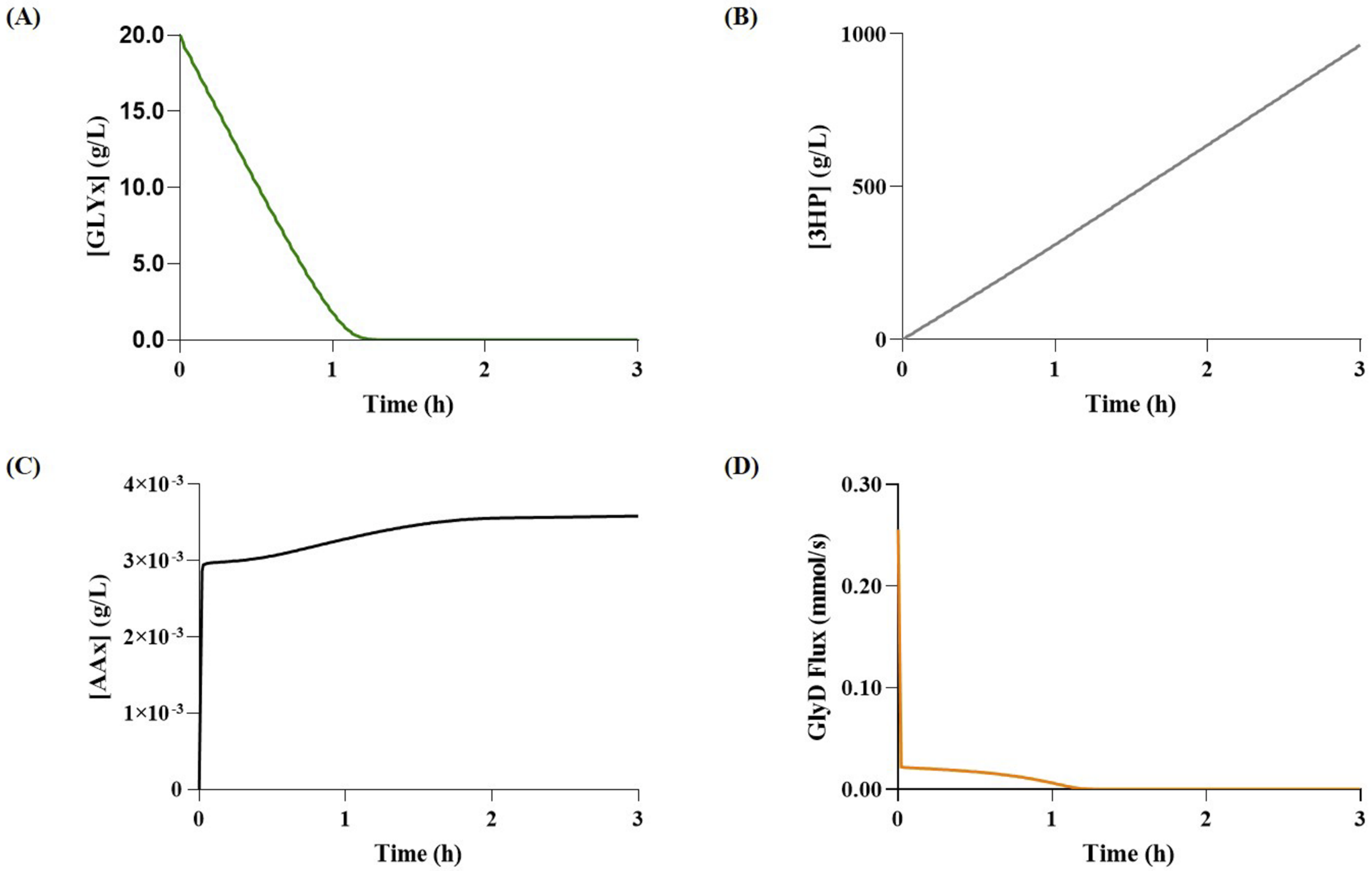

(A)
[GLYx] (g/L)
20.0
15.0
10.0
5.0
0.0
0
1
2
3
Time (h)
(B)
[3HP] (g/L)
1000
500
0
0
1
2
3
Time (h)
(C)
[AAx] (g/L)
4×10-3
3×10-3
2×10-3
1×10-3
0
0
1
2
3
Time (h)
(D)
GlyD Flux (mmol/s)
0.30
0.20
0.10
0.00
0
1
2
3
Time (h)

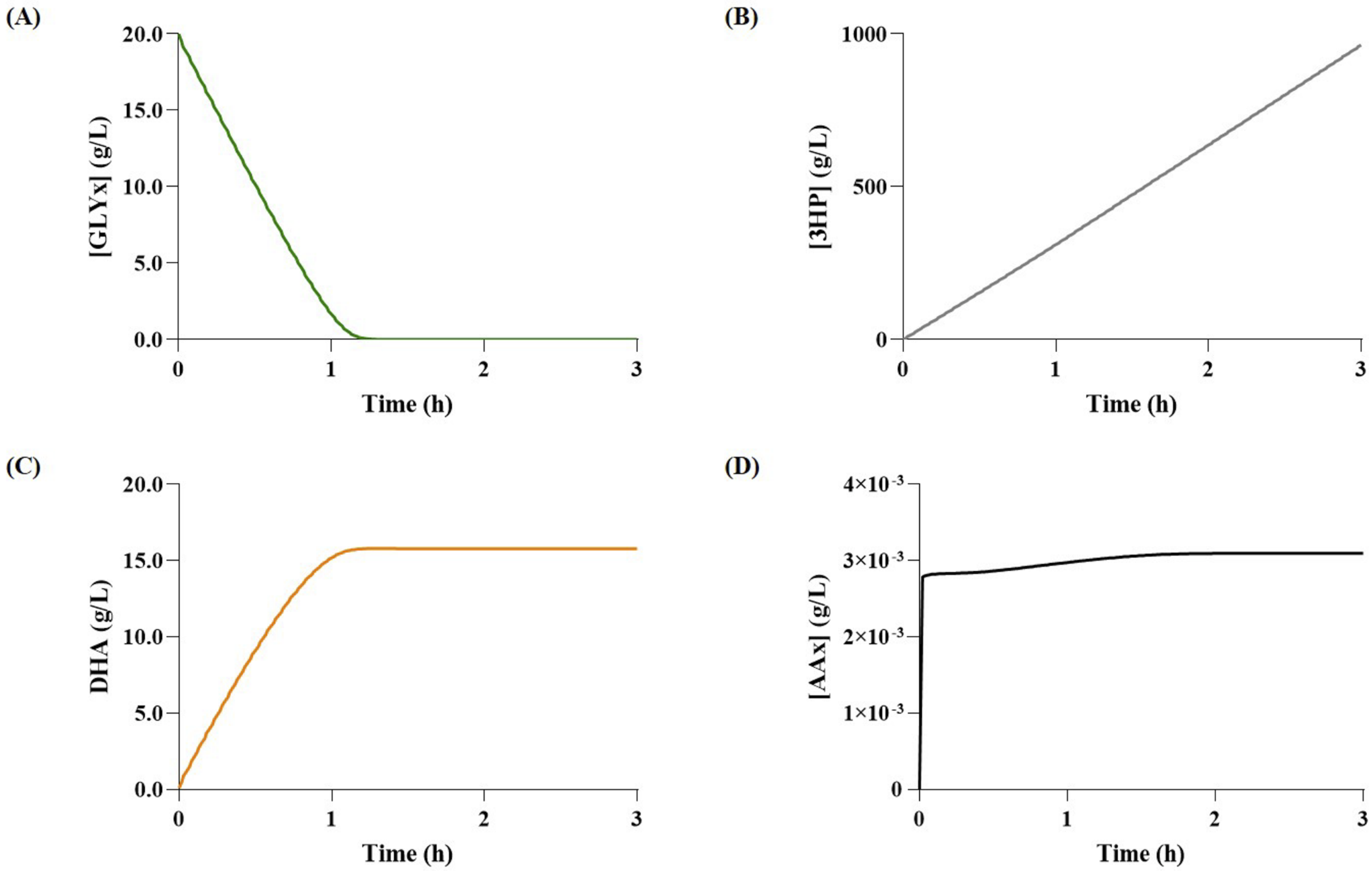

(A)
[GLYx] (g/L)
20.0
15.0
10.0
5.0
0.0
0
1
2
3
Time (h)
(B)
[3HP] (g/L)
1000
500
0
0
1
2
3
Time (h)
(C)
DHA (g/L)
20.0
15.0
10.0
5.0
0.0
0
1
2
3
Time (h)
(D)
[AAx] (g/L)
4×10-3
3×10-3
2×10-3
1×10-3
0
0
1
2
3
Time (h)

### S2 Appendix

#### 1 Kinetic Data

In this section it is laid out the kinetic information found in a variety of databases for all the reactions that needed to be included in the model. First are described the reactions for the central carbon metabolism (CCM) extension, then the heterologous reactions required to produce acrylic acid (AA), and finally the transport reactions added to transport end metabolites to the extracellular compartment.

##### 1.1 Extension of the Central Carbon Metabolism (CCM)

The first step to produce AA is to extend the CCM to include the production of glycerol, malonyl-CoA, and *β*-alanine, which are crucial intermediary metabolites for the heterologous pathways and naturally produced in *Escherichia coli*. Nine reactions were required to obtain those metabolites. For the glycerol case, two enzymes allow to produce glycerol and are catalyzed by the glycerol-3-phosphate dehydrogenase (G3pD) and the glycerol-3-phosphate phosphatase (G3pP), and three allow the utilization of glycerol as a carbon source, catalyzed by the glycerol kinase (GlyK), glycerol dehydrogenase (GlyD) and the dihydroxyacetone phosphate transferase (DhaPT). To produce malonyl-CoA, only one reaction was added, and it is catalyzed by the acetyl-CoA carboxylase (AccC). Finally, the remaining three reactions, catalyzed by the aspartate aminotransferase (AspAT), the aspartate carboxylase (AspC), and the *L*-glutamate dehydrogenase (GluD), are required to produce *β*-alanine.

###### 1.1.1 *Glycerol-3-phosphate dehydrogenase (G3pD)*

This enzyme is responsible for the conversion of dihydroxyacetone phosphate to glycerol-3-phosphate, coupled with the oxidation of NADPH to $NADP^+$. Three works were found regarding this enzyme [29,30,31] (Kito and Pizer, 1969; Edgar and Bell, 1978b, Edgar and Bell, 1978a) The kinetic properties of G3pD were fully characterized in the work of Edgar and Bell (1978) [21]. They reported a $K_m$ of 0.18 mM for dihydroxyacetone phosphate ($K_{m,a}$), 0.0034 mM for NADPH ($K_{m,b}$), 0.03 mM for glycerol-3-phosphate ($K_{m,p}$), and finally 0.165 mM for $NADP^+$ ($K_{m,q}$) and a $K_{eq}$ of 900 for this reaction. They did not accurately identify the kinetic mechanism of the enzyme, however they stated that the most probable one was the Rapid Equilibrium Random Bi Bi. Later that year, the same authors published another work concerning the kinetic properties of this enzyme where they reported two more $K_m$ values for NADPH, 0.0041 mM and 0.0037 mM [30]. Moreover, a characterization made by Kito and Pizer (1969) [31] reported $K_m$ values for dihydroxyacetone phosphate and glycerol-3-phosphate of 0.17 mM and 0.21 mM, respectively. This combination of parameters was used to calculate the mean values that were reported in Table 1.

##### 1.1.2 *Glycerol-3-phosphate phosphatase (G3pP)*

The G3pP is responsible for the production of glycerol from glycerol-3-phosphate. For this enzyme, only the work of Salles et al. (2007) [32] was considered. This enzyme follows a single substrate Michaelis-Menten kinetic, and according to their work, the respective $K_m$ for glycerol-3-phosphate was equal to 2.9 mM, as shown in Table 1.

##### 1.1.3 *Glycerol-3-phosphate phosphatase (G3pP)*

This enzyme uses the dephosphorylation of ATP to convert glycerol into glycerol-3-phosphate. The kinetic parameters of this enzyme were found in different works. Pettigrew et al. (1990) [33] reported a $K_m$ for glycerol ($K_{m,a}$) of 0.0049 mM and for ATP ($K_{m,b}$) of 0.0084 mM, and a dissociation constant for ATP ($K_{d,a}$) of 0.086 mM, and they also reported that the enzyme seemed to have a random Bi Bi kinetic mechanism. In turn, Hayashi and Lin (1967) [34] reported a $K_{m,a}$ of 0.0013 mM, Thorner and

Paulus (1973) [35] reported a $K_{m,a}$ of 0.01 mM, and finally, the work from Applebee et al. (2011) [36] reported a $K_{m,b}$ of 0.0078 mM. The resulting mean values for each parameter are presented in Table 1.

##### 1.1.4 *Glycerol dehydrogenase (GlyD)*

The GlyD is the enzyme responsible for the conversion of glycerol into dihydroxyacetone, coupled with the reduction of $NAD^+$ to NADH. This reaction was characterized in *E. coli* by Piattoni et al. (2013) [51]. In their work they presented six different $K_m$ values for Glycerol ($K_{m,a}$), 76, 81,47, 46, 13 and 24 mM, and for $NAD^+$ ($K_{m,b}$), 0.81, 1, 3.2, 0.8, 1.4 and 1.1 mM. Moreover, according to the authors, this enzyme exhibited a behavior that seemed to fit hill cooperativity kinetics, and thus they also proceeded to report six values for the hill coefficient ($n$) of 0.9, 0.9, 1, 1,1.1 and 1. The resulting mean values for each parameter are presented in Table 1.

##### 1.1.5 *Dihydroxyacetone phosphate transferase (DhaPT)*

The DhaPT enzyme catalyzes the final reaction that connects glycerol to the remain of the CCM, through the conversion of dihydroxyacetone into dihydroxyacetone phosphate, an intermediary of the glycolysis pathway, coupled with the dephosphorylation of phosphoenolpyruvate to pyruvate. This enzyme was not fully characterized for both substrates, as the literature revealed a lack of $K_m$ values for phosphoenolpyruvate in the used databases. Therefore, mass action kinetics were used to represent the dynamics of this reaction, and the $k$ value was calculated using method 1 (S2 Table 2).

##### 1.1.6 *Acetyl-CoA carboxylase (AccC)*

This enzyme is responsible for the conversion of acetyl-CoA to malonyl-CoA coupled with the dephosphorylation of ATP. The kinetic parameters of this enzyme were described by Soriano et al. (2005) [37], in which they reported a $K_m$ for acetyl-CoA ($K_{m,a}$) of 0.018 mM and for ATP ($K_{m,b}$) of 0.06 mM. Moreover, a study by Freiberg et al. (2004) [38] and another by Meades et al. (2010) [39], showed that malonyl-CoA, a product of this reaction, presented competitive inhibition towards acetyl-CoA, and therefore both presented values for the inhibition constant for this metabolite ($K_{i,p}$) of 0.1 mM and 0.04 mM, respectively. These parameters were used to calculate the mean values reported in Table 1.

##### 1.1.7 *Aspartate aminotransferase (AspAT)*

The AspAT is responsible for the formation of *L*-aspartate and $\alpha$-ketoglutarate from oxaloacetate and *L*-glutamate. Yagi et al. (1985) [44] presented a kinetic description of this enzyme in which $K_m$ values of 15 mM, 0.01 mM, 0.24 mM, and 1.3 mM, were reported for *L*-glutamate ($K_{m,a}$), oxaloacetate ($K_{m,b}$), aspartate ($K_{m,p}$), and $\alpha$-ketoglutarate ($K_{m,q}$), respectively. In addition, $K_m$ values were also reported by Chow et al. (2004) [45] for $\alpha$-ketoglutarate of 0.48 mM, by Mavrides and Orr (1975) [46] of 0.37 mM and 4.4 mM for oxaloacetate and *L*-aspartate respectively, by Deu et al. (2002) [47] for $\alpha$-ketoglutarate of 0.59 mM and aspartate of 1.9 mM, and finally, by Fernandez et al. (2012) [48] of 37 mM and 5.2 mM for *L*-glutamate and for *L*-aspartate of 3.1 and 4 mM. Unfortunately, the underlying mechanism of this enzyme was not specified in this paper, but according to BioCyc, this enzyme is known to follow a reversible Ping-Pong Bi Bi mechanism. Moreover, to finish this enzyme characterization, the eQuilibrator database was accessed to retrieve the $K_{eq}$, whose value was 3.2. The mean values are presented in Table 1.

##### 1.1.8 *Aspartate carboxylase (AspC)*

This enzyme catalyzes the conversion of *L*-aspartate to $\beta$-alanine. Like the G3pP, this enzyme is characterized by a single substrate Michaelis-Menten kinetic with a $K_m$ of 0.151 mM accordingly to Ramjee et al. (1997). The same mechanism was described by Williamson and Brown (1979) [50], but with a similar $K_m$ value of 0.16 mM, which resulted in a mean value of 0.156 mM (Table 1).

##### 1.1.9 *L-glutamate dehydrogenase (GluD)*

The reaction catalyzed by GluD was included because the dynamic model used did not include *L*-glutamate, which is required by AspAT. GluD is responsible for the conversion of the $\alpha$-ketoglutarate to *L*-glutamate, coupled with the oxidation of NADPH to $NADP^+$. The used kinetic parameters for this enzyme were found in four different works. Considering the $K_m$ value for $\alpha$-ketoglutarate ($K_{m,a}$) and $NADP^+$ ($K_{m,b}$), the reported values were, respectively, 0.68 mM and 0.0597 mM according to Sharkey and Engel (2008) [40], 0.64 mM and 0.040 mM by Sakamoto et al. (1975) [41], 0.46 mM and 0.012 mM by Mäntsälä and Zalkin (1976) [42], and finally, 0.2 mM and 0.035 mM by Di Fraia et al. (2000) [43]. In addition, the kinetic mechanism assumed was a simple Michaelis-Menten kinetics. These parameters were used to calculate the mean values that are reported in Table 1.

#### 1.2 Pathways for Acrylic Acid Production

Three different heterologous pathways were identified to produce AA, the glycerol, malonyl-CoA, and $\beta$-alanine routes. These pathways differ from each in the production of 3-hydroxypropionate (3-HP),. The glycerol pathway includes two reactions to produce 3-HP, which are catalyzed by the glycerol dehydratase (GlyDH) and the 3-hydroxypropionaldehyde dehydrogenase (3hpaD). Regarding malonyl-CoA pathway, two reactions are needed to produce 3-HP, namely the ones catalyzed by the malonyl-CoA reductase (McoaR) and the malonic semialdehyde reductase (MsaR). And finally, two for the $\beta$-alanine pathway, $\beta$-alanine aminotransferase (BaAT) and the MsaR are needed to convert $\beta$-alanine to 3-HP. After obtaining 3-HP, three more enzymes are required, the 3-hydroxypropionyl-CoA synthase (3hpcoaS), 3-hydroxypropionyl-CoA dehydratase (3hpcoaDH), and the acrylyl-CoA thioesterase (AcoaTioE).

##### 1.2.1 *Glycerol dehydratase (GlyDH)*

This enzyme is responsible for the conversion of glycerol to 3-hydroxypropionaldehyde (3-HPA). The kinetic parameters of the enzyme in *Lactobacillus collinoides* were determined by Sauvageot et al. (2002) [53]. In their work they reported a $K_m$ of 8.3 mM for glycerol, and an activation constant ($K_a$) of 0.008 mM for vitamin $B_{12}$. Moreover, they also reported a specific activity of 0.018 $mmol.min^{-1}.mg^{-1}$ when glycerol was the substrate, and a molecular weight of 207 kDa, which allowed the calculation of a $K_{cat}$ value of 0.0621 $s^{-1}$. In addition, a $K_m$ value for glycerol (4.0 mM) reported by Schütz and Radler (1984) [54] in *Lactobacillus brevis* was also considered for the mean calculation because of the taxonomy proximity between both species. Thus, considering the available kinetic parameters, and the fact that vitamin $B_{12}$ is an activator of the reaction and not an intermediary, the specific activation mechanism rate law was assumed (Table 2).

##### 1.2.2 *3-hydroxypropionaldehyde dehydrogenase (3hpaD)*

The following step of the heterologous pathway is catalyzed by the 3hpaD, and it consists on the conversion of 3-HPA to 3-hydroxypropionate (3-HP) coupled with the reduction of $NAD^+$ to NADH. The kinetic properties of the 3hpaD from *E. coli* were studied by Jo et al. (2008) [55]. This study reported a $K_{cat}$ of 28.54 $s^{-1}$ when 3-HPA was used as a substrate, as well as $K_m$ values of 0.49 mM and 0.06 mM for 3-HPA ($K_{m,a}$) and $NAD^+$ ($K_{m,b}$), respectively. Considering that no kinetic mechanism was associated with this enzyme, a simple two substrate Michalis-Menten equation was assumed (Table 2). Furthermore, since this reaction had no flux in the stoichiometric model when using Method 1, the $V_{max}$ was calculated using the method for the heterologous reactions (Method 2).

##### 1.2.3 *Malonyl-CoA reductase (McoaR)*

The McoaR is responsible for the first reaction on the malonyl-CoA pathway, which consists in the formation of malonic semialdehyde (MSA) from malonyl-CoA, coupled with the oxidation of NADPH to $NADP^+$ and the release of a coenzyme A molecule. This enzyme is present in *Chloroflexus aurantiacus* and was described by Hügler et al. (2002) [56], which reported a $K_{cat}$ of 50 $s^{-1}$ when malonyl-CoA was used as a substrate, and $K_m$ values of 0.3 mM and 0.025 mM for malonyl-CoA ($K_{m,a}$) and NADPH ($K_{m,b}$), respectively. The authors also stated that the enzyme exhibited a two substrate Michaelis-Menten behavior (Table 2). Furthermore, the values reported by Liu et al. (2013) [57] were also accounted to determine the mean values of each parameter. They reported two $K_m$ values for NADPH of 0.0238 and 0.0417 mM.

##### 1.2.4 *Malonic semialdehyde reductase (MsaR)*

The MsaR enzyme, which catalyzes the final reaction of the malonyl-CoA pathway, was described by Kockelkorn and Fuchs (2009) [58]. This enzyme is responsible for the conversion of MSA to 3-HP, coupled with the oxidation of NADPH to $NADP^+$. The authors characterized this enzyme from *Metallosphaera sedula* and were able to identify its kinetic parameters. The $K_{cat}$ value reported when malonyl-CoA was used as the substrate was 115 $s^{-1}$, and the $K_m$ values were 0.07 mM for both MSA ($K_{m,a}$) and NADPH ($K_{m,b}$). Finally, a two substrate Michaelis-Menten mechanism was assumed for this model (Table 2).

##### 1.2.5 *β-alanine aminotransferase (BaAT)*

This enzyme catalyzes the conversion of *β*-alanine and *α*-ketoglutarate to MSA and *L*-glutamate. Furthermore, *E. coli*'s BaAT was described by Liu et al. (2005) [59]. The authors believed that the enzyme followed Ping-Pong Bi Bi mechanism with competitive substrate inhibition for *α*-ketoglutarate and proceeded to estimate the respective parameters. They reported a $K_{cat}$ of 47.4 $s^{-1}$, and a $K_m$ for $\beta$-alanine ($K_{m,a}$) and $\alpha$-ketoglutarate ($K_{m,b}$), of 5.8 mM and 1.07 mM, respectively. Furthermore, the $K_{i,b}$ for $\alpha$-ketoglutarate was also estimated, and was equal to 10.2 mM (Table 2). Even though this enzyme was characterized from *E. coli*, it was not present in the stoichiometric model, thus the $V_{max}$ was calculated using the Method 2.

##### 1.2.6 *3-hydroxypropionyl-CoA synthase (3hpcoaS)*

This enzyme uses the dephosphorylation of ATP to join a coenzyme A molecule to 3-HP, thus forming 3-hydroxypropionyl-CoA (3-HP-CoA). Alber and Fuchs (2002) [60] studied a variant of the 3hpcoaS from *Chloroflexus aurantiacus*. In their work they stated that the enzyme probably follows a Michaelis-Menten mechanism, and the parameters for that mechanism were a $K_{cat}$ of 36 $s^{-1}$, and the $K_m$ values for 3-HP ($K_{m,a}$) of 0.015 mM, CoA ($K_{m,b}$) of 0.01 mM, and ATP ($K_{m,c}$) of 0.05 mM (Table 2).

##### 1.2.7 *3-hydroxypropionyl-CoA dehydratase (3hpcoaDH)*

The following reaction, which is catalyzed by the 3hpcoaDH, is responsible for the formation of acrylyl-CoA (AA-CoA) from 3-HP-CoA. A variant of the 3hpcoaDH from *M. sedula* was characterized by Teufel et al. (2009) [61], whose work helped to understand the underlying mechanism that controls the enzyme's activity. According to this study, the enzyme follows a Michaelis-Menten kinetics, with a $K_{cat}$ of 96 $s^{-1}$, and $K_m$ value of 0.06 mM for 3-HP-CoA (Table 2).

##### 1.2.8 *Acrylyl-CoA thioesterase (AcoaTioE)*

The final reaction consists in the conversion of AA-CoA to AA coupled with the release of a coenzyme A molecule. No studies characterizing this enzyme's kinetic parameters were found in the literature. Hence, an enzyme catalyzing a similar reaction was sought. Ultimately, *E. coli*'s acyl-CoA thioesterase was selected as surrogate. This enzyme was the one used in the work of Chu et al. (2015)

[1], which using a direct bio-based route achieved AA production for the first time. As opposed to the original reaction, this one was characterized in a report by Zhuang et al. (2008) [62], where it was reported that this enzyme kinetics follows the Michaelis-Menten model, with a $K_{cat}$ of 96 $s^{-1}$, and $K_m$ for AA-CoA of 0.167 mM (Table 2).

##### 1.3 Transport Reactions

The exchange reactions allow the transport of 3-HP (XCH_3HP), AA (XCH_AA), glycerol (XCH_GLY), malonyl-CoA (XCH_MCOA), and *β*-alanine (XCH_BA) from the cytoplasm to the extracellular compartment and vice-versa. Since no kinetic data concerning transport reactions for these metabolites was found or even evidence of the existence of such mechanisms in the case of 3-HP and AA, these reactions were included akin to the ones in the original model [19]. Furthermore, this entire set of reactions follows the reversible Michaelis-Menten kinetics, with a $V_{max}$ of 100 mM/s and a $K_m$ of 10 mM, as it was arbitrarily chosen in the original model [19].
